# The *Mosigvirus* AV110 Elongasome-Modulating Peptide (Emp) inhibits *Escherichia coli* growth by targeting Mre-dependent peptidoglycan synthesis

**DOI:** 10.64898/2026.09.14.751387

**Authors:** Saar S. F. Van Overfelt, Anders Nørgaard Sørensen, Viktor Hundtofte Mebus, Jakob Rask Tornby, Jonathan Brewer, Rob Lavigne, Lone Brøndsted

## Abstract

Extended-spectrum β-lactamase and AmpC-producing *Escherichia coli* (ESBL/AmpC *E. coli*) are increasingly recognized as public health threats due to their resistance to commonly used antibiotics. With limited treatment options available, new antibacterials and antibacterial targets can be explored by studying early-expressed phage proteins involved in host reprogramming, as these proteins interact with and disrupt essential bacterial processes. A systematic screening of 45 early-expressed proteins with unknown function of *Mosigvirus* AV110 identified six proteins that inhibit the growth of ESBL/AmpC *E. coli*, suggesting a role in host reprogramming. In-depth analysis of Gp94, termed Elongasome-Modulating Peptide (Emp), demonstrated that it alters cell morphology and the localization of peptidoglycan synthesis. This effect could be partially complemented by MreB and MreD. Since Emp interacts directly with MreB, MreC, and MreD, we propose that Emp perturbs elongasome-associated peptidoglycan synthesis, thus providing a scaffold for developing new antibacterials against ESBL/AmpC-producing *E. coli*.

## Introduction

*Escherichia coli* is a genetically highly diverse species, comprising pathogenic and commensal strains, both potentially encoding various antibiotic resistance genes. Those resistance genes often include plasmid-encoded extended-spectrum β-lactamase (ESBL) and chromosomal *ampC* genes. ESBL/AmpC *E. coli* are commensals in food-producing animals, including pigs, bovines, broilers, and turkeys, and are frequently detected in the meat of these animals^1^. These strains are highly diverse and belong to all known *E. coli* phylogroups^2,3^, with a significant variation in the ESBL/AmpC genes, with CTX-M-1 and CMY-2 types being the most prevalent^2,3^. These ESBL genes are often carried on conjugative plasmids, enabling horizontal transfer to pathogenic strains. However, their importance and impact on these processes are not fully understood^2,3^. In addition to food-producing animals, humans can be asymptomatic carriers of ESBL/AmpC *E. coli*^4^. Moreover, this bacterium also causes community- and hospital-acquired infections^5^, with an increase in community-onset infections between 2007 and 2017 in countries like Denmark^6^. The resistance of ESBL/AmpC *E. coli* to penicillin, aztreonam, and first-, second-, and third-generation cephalosporins, often accompanied by cross-resistance to other antibiotic classes, severely limits therapeutic options and poses a major public health challenge^5–8^.

Phages, bacterial viruses, have been proposed as an alternative to antibiotics for treating multidrug-resistant infections. In clinical settings, phage therapy is often combined with antibiotics to promote clinical improvement and, in some cases, completely eradicate *E. coli*^9–11^. However, a combination strategy is not always efficient when targeting multidrug-resistant bacteria^11,12^. Therefore, alternative approaches are needed, such as inhibiting bacterial growth by phage proteins involved in reprogramming the bacterial metabolism towards phage production^13^. Bacterial reprogramming is achieved by phage proteins expressed during the early stage of infection that interfere with the host transcriptome, proteome, and metabolome. For example, in *E. coli*, host RNA transcripts are rapidly depleted during a phage T4 infection and replaced with phage RNA transcripts, indicating an almost complete shutdown and degradation of the host transcriptome^14^. In contrast to the transcriptome, the *E. coli* proteome remains constant during T4 infection, indicating that there is no enhanced degradation of host proteins^14^. However, host proteins can be repurposed to assist in the progression of the phage infection. This is observed with the RNA polymerase, which is redirected from transcribing host genes to transcribe the T4 early genes because of an ADP-ribosylation of the polymerase by the T4 protein Alt co-injected with the genome and the stronger promoter of the T4 early genes^15^. Lastly, the metabolome is also influenced by this host reprogramming, although the effect is highly phage-specific^16^.

Early-expressed proteins involved in bacterial reprogramming have been proposed as potential antibacterials due to their ability to interfere with essential processes and cause growth inhibition^17,18^. Examples include the *Tequatrovirus* T4 protein Alc, the *Tequintavirus* T5 protein 008 (Hdi), *Teseptimavirus* T7 protein gp0.6, and the *Bruynoghevirus* LUZ24 protein gp9 (Igy)^19–21^. Alc induces site-specific transcription termination by binding selectively to the cytosine-containing host DNA^19^. This activity results in the shutdown of host transcription while preserving the T4 transcription. The T5 protein Hdi inhibits the growth of *E. coli* by interfering with cell division^20^. This occurs through interaction with the cell-division protein FtsZ, conferring a competitive advantage to T5 phages with an active Hdi over Hdi-mutant. Additionally, Gp0.6 of phage T7 interacts with MreB, thereby inhibiting the formation of the elongasome required for maintaining a rod-shaped morphology during cell division^22^. Lastly, the LUZ24 protein Igy inhibits the enzymatic activity of *Pseudomonas* DNA gyrase, resulting in filamentous growth, although its biological function during phage infection remains unclear^21^. Despite the increasing number of early-expressed proteins being characterized, the functions and targets of many early proteins remain unknown. These uncharacterized proteins hold potential for discovering novel ways in which phages reprogram their host, which could lead to identifying novel antibacterial targets^23^.

An extensive collection of ESBL-targeting phages, including ten phages belonging to the *Mosigvirus* genus, was recently isolated and characterized^24^. Mosigviruses are members of the *Tevenvirinae* subfamily and can be isolated from diverse environments, for example, wastewater and animal feces^24,25^. Their host range varies from small to moderately broad^24–26^, and they are generally resistant to DNA-targeting defense systems including restriction-modification systems^27^. Like other *Tevenvirinae* phages, they have a relatively large genome of approximately 170 kb that encodes numerous proteins of unknown function^28^. In contrast to the well-studied *Tequatrovirus* phages, the specific proteins involved in host reprogramming by *Mosigvirus* phages, as well as the molecular mechanisms, remain poorly understood.

Our first objective was to identify the early-expressed proteins of the *Mosigvirus* phage AV110, which was previously isolated from pig waste and infects a diverse range of ESBL/AmpC *E. coli*^24^. Next, we aimed to identify the early-expressed proteins that inhibit the growth of ESBL/AmpC *E. coli*. In addition, we sought to elucidate the mode of action of one of the early-expressed proteins, Gp94 (Emp), and discovered that it interferes with the localization of peptidoglycan synthesis. Lastly, we determined that Emp interacts with the elongasome proteins MreB and MreD. Taken together, these results suggest that Emp targets MreB-dependent peptidoglycan synthesis, offering a new strategy for developing novel antimicrobials to combat antibiotic-resistant bacteria.

## Results

### Identification of early proteins in phage AV110

Phages AV110 and T4 are classified within the subfamily *Tevenvirinae* but belong to distinct genera, i.e. *Mosigvirus* and *Tequatrovirus*, respectively. They share an overall nucleotide identity of 61% and contain many homologous genes organized similarly in their genomes (Figure 1A). This genomic syntheny between AV110 and T4 was used to predict putative early-expressed proteins of phage AV110. Ninety early-expressed proteins of T4 were recently identified in a time-resolved transcriptome and proteome analysis of T4 infecting *E. coli*^14^. The amino acid sequences of these 90 early-expressed T4 proteins were extracted and aligned with 269 CDS of AV110. Proteins of AV110 that aligned with an early protein of T4 were identified as a putative early protein, resulting in the identification of 63 putative early-expressed proteins spread over the genome (Figure 1A and **Error! Reference source not found.**). Based on a literature review of the T4 early proteins, 18 genes of known function or known growth-inhibitory effect on their host were excluded from our selection (**Error! Reference source not found.**). This led to the identification of 45 putative early-expressed proteins encoded by phage AV110 with unknown or poorly characterized functions (Figure 1B).

**Figure 1:**
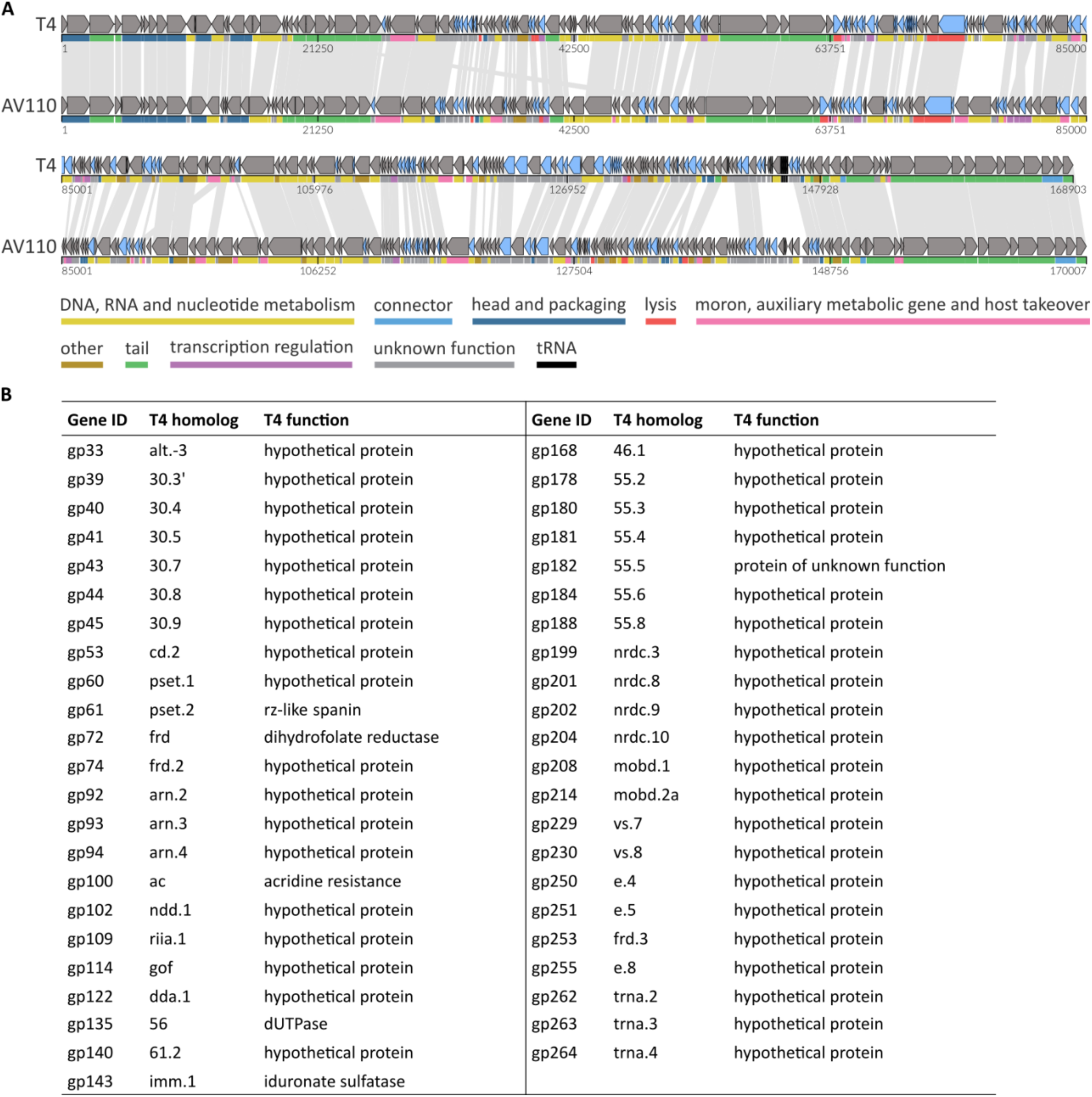
Early proteins of AV110 are identified based on sequence similarity with T4. A) A whole-genome alignment of T4 and AV110 was created and visualized using LoVis4u^29^ with indication of the homologous regions. The functional category of each gene is indicated below the gene. Early-expressed proteins of T4^14^, and their homologs in AV110, which were identified through sequence alignment, are indicated in blue. B) Overview of the 45 identified early-expressed proteins in AV110 and their T4 homolog with an unknown or poorly characterized function. See also **Error! Reference source not found.**.

### Early proteins impact the growth of *E. coli* ESBL098

To evaluate the impact on growth of expressing the 45 putative early-expressed proteins of AV110, we used *E. coli* ESBL098. This strain was previously isolated from broiler meat and belongs to phylogroup B1 and sequence type ST-1431^2^. It encodes the ESBL gene *bla*_CTX-M-1_ on an IncI1-type plasmid and has additional resistance to fluoroquinolones and sulfonamides.

To assess the growth-inhibitory effect, each early gene was cloned into the low-copy number vector pPS26 under the control of the rhamnose-inducible promoter *p_rhaB_*. After the transformation of ESBL098, growth was evaluated by spotting tenfold dilutions on LA plates with and without rhamnose. The empty vector and the T4 gene *alc*, which is known to terminate transcription in *E. coli^19^*, served as the negative and positive control, respectively. The empty vector did not affect the growth of ESBL098, while induction of *alc* resulted in a four-log reduction in growth (Figure 2). Of the 45 tested genes, the expression of six genes affected the growth of ESBL098 (Figure 2 and **Error! Reference source not found.**). Gp135 and Gp204 were identified as growth-inhibitory proteins and are described elsewhere^30^. Expression of *gp94* showed the most substantial impact on growth, resulting in an approximate three-log reduction. Smaller colonies could be observed at higher dilutions, suggesting selection of mutants that counter the effect of the phage proteins or prevent *gp94* expression. Upon expressing *gp208*, *gp214*, and *gp263*, growth was still observed in all dilutions, although the spots were less densely grown compared to the non-induced control. This could potentially be due to the metabolic burden of overexpressing a foreign gene, rather than a specific effect of the protein itself. Also, smaller colonies that might have obtained resistance against Gp208, Gp214, or Gp263 were observed. These results show that the early-expressed proteins Gp94, Gp208, Gp214, and Gp263 of AV110 inhibit the growth of ESBL098 to varying levels, highlighting their potential for discovery of novel antibacterial targets.

**Figure 2:**
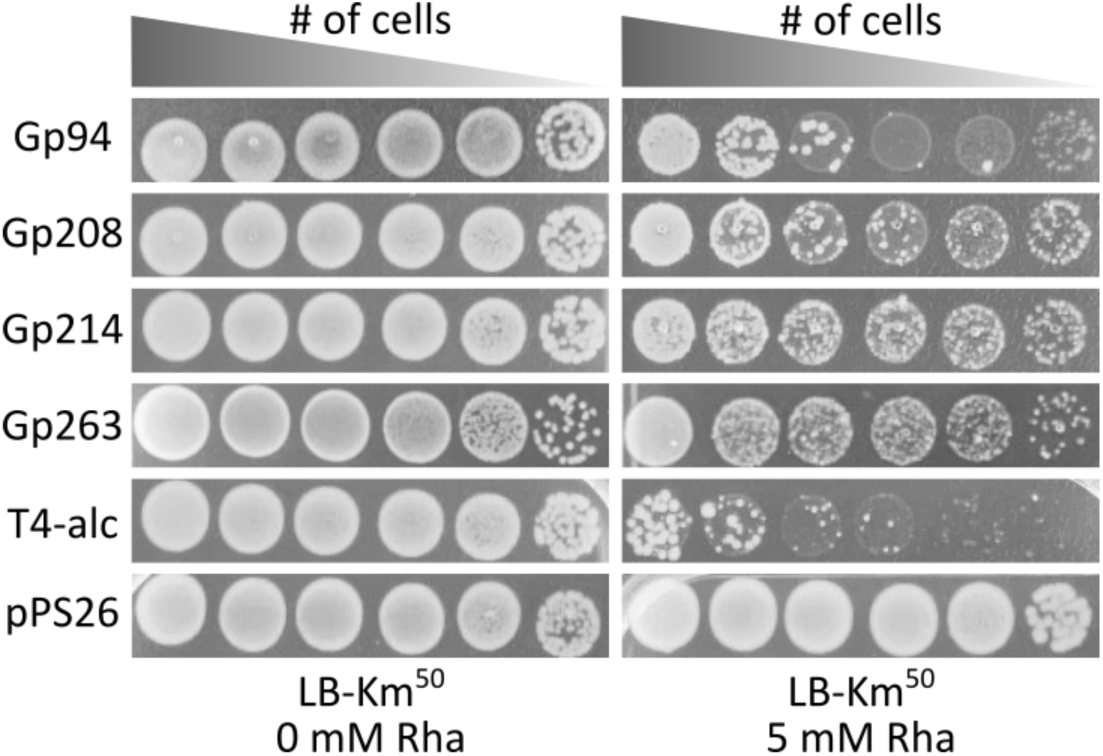
Four early phage proteins influence the growth of ESBL098. Tenfold dilutions of ESBL098 transformed with the plasmid pPS26 encoding the respective phage gene were spotted on LA plates with and without rhamnose to induce the expression of the phage protein. The T4 gene *alc* and the empty vector pPS26 served as the positive and negative controls. See also **Error! Reference source not found.**.

### Early proteins target diverse processes in ESBL *E. coli*

To investigate whether Gp94, Gp208, Gp214, and Gp263 affect the growth of a broader range of *E. coli* strains, individual genes were expressed in seven additional ESBL/AmpC *E. coli* strains from different phylogroups and sequence types and *E. coli* Stellar™. As before, the growth-inhibitory effect was evaluated by spotting a tenfold dilution series on LA plates with and without rhamnose. Gp94 inhibited the growth of all eight strains, ranging from complete inhibition in ESBL087 to an approximate four-log reduction in Stellar^TM^ (Figure 3A). Gp208 affected ESBL049 but not the closely related ESBL032 or any of the other tested strains (Figure 3B). Gp214 and Gp263 did not inhibit the growth of any of the additional ESBL/AmpC *E. coli* strains or Stellar^TM^ (Figure 3C-D). These results indicate that Gp94 inhibits a conserved and essential process in *E. coli*, whereas Gp208, Gp214, and Gp263 interfere with a strain-specific target.

**Figure 3:**
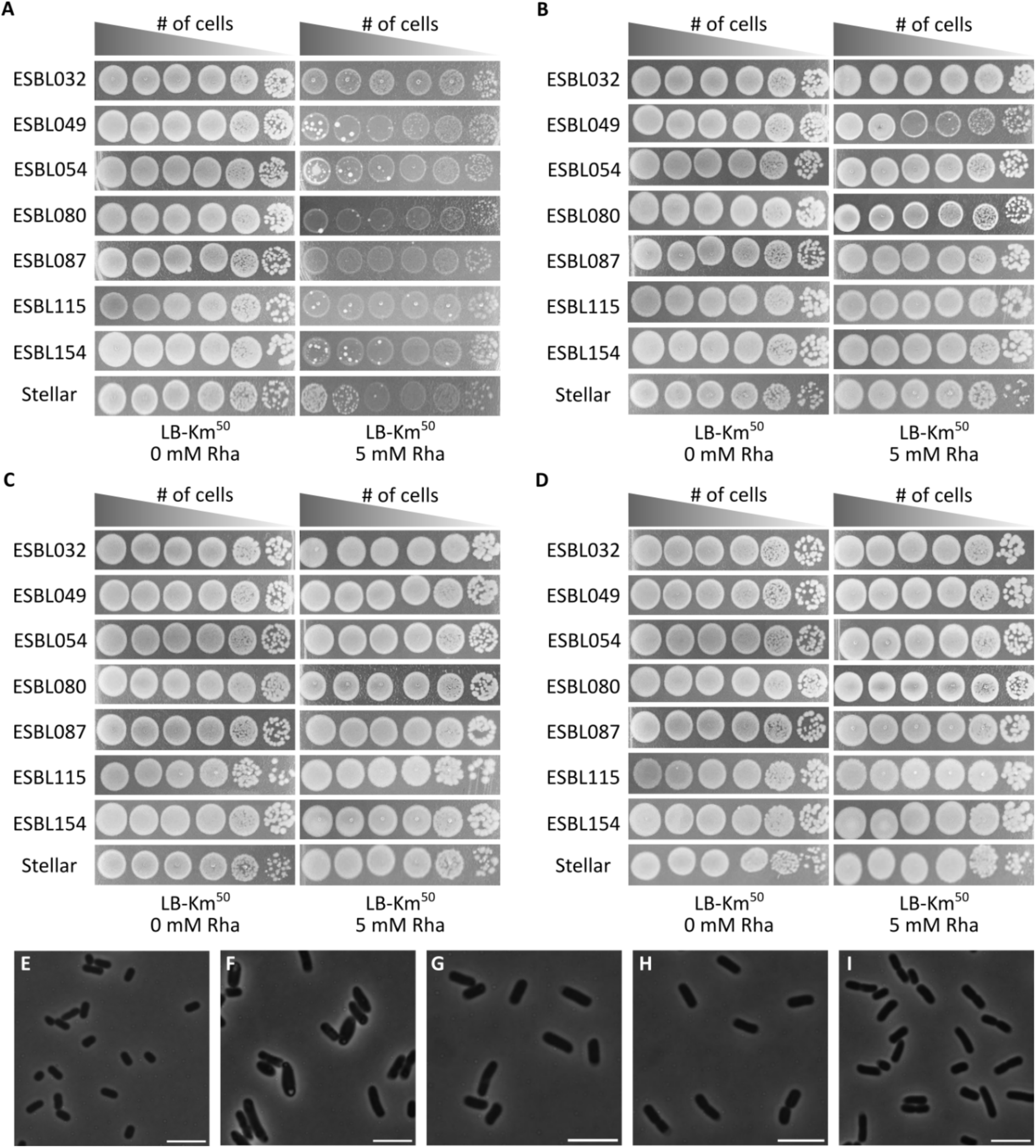
Early-expressed proteins have diverse effects on ESBL/AmpC *E. coli.* Tenfold dilutions of diverse ESBL/AmpC *E. coli* strains and Stellar^TM^, transformed with A) pPS26::*gp94*, B) pPS26::*gp208*, C) pPS26::*gp214*, and D) pPS26::*gp263* were spotted on LA plates without and with rhamnose to assess the growth-inhibitory effect. Phase contrast microscopy after four hours of induction of ESBL098 transformed with E) the empty vector pPS26, F) pPS26::*gp94*, G) pPS26::*gp208*, H) pPS26::*gp214*, and I) pPS26::*gp263*. Representative images out of 15 images for each condition are shown. Scale bar = 5 µm.

Next, the effect of the growth-inhibitory phage proteins on the morphology of ESBL098 was assessed using phase-contrast microscopy. After four hours of inducing expression, Gp94, Gp208, Gp214, and Gp263 resulted in cells that were slightly larger than the control (p < 0.001). This cell elongation is likely due to the growth-inhibitory effect of each protein, which also prevents cells from reaching the stationary phase, during which cells usually become shorter. Gp94 expression resulted in the longest cells, with an average length of 4.05 ± 1.22 µm, compared to 1.85 ± 0.41 µm in the control. On addition, Gp94 expression resulted in cell bloating, either at the cell poles or in the middle of the cell, whereas expression of the other proteins did not affect cell shape (Figure 3E-I). Since Gp94 causes a significant growth-inhibitory effect in a diverse set of ESBL/AmpC *E. coli* and a striking cell morphology change in ESBL098, the remainder of this study will focus on this protein.

### Gp94 is conserved across *Tevenvirinae* and exhibits a protein fold of unknown function

To better understand the properties and potential role of the 109 amino acid protein (12.74 Kda) Gp94 (later termed Emp), we carried out *in silico* analyses, including gene conservation and structural predictions. Gp94 is conserved within *Mosigvirus* phages, but it is also encoded by phages belonging to the *Dhakavirus, Gaprivervirus, Kanagawavirus, and Tequatrovirus* genera within *Tevenvirinae*. On the AV110 phage genome, Gp94 is flanked by three other putative early-expressed proteins. Two of these genes, *gp92* and *gp93*, are of unknown function, while the third gene, *gp95*, encodes a MotA-like transcriptional regulator of middle promoters. However, Gp94 was the only gene in this region with a growth-inhibitory effect on ESBL098, although Gp95 was not included since its function was already established (**Error! Reference source not found.**). To better understand the function of Gp94, its protein structure was confidently predicted by AlphaFold 3^31^, demonstrating an N-terminal alpha helix, followed by five anti-parallel beta strands and two alpha helices connected by a flexible region (**Error! Reference source not found.**A). Furthermore, the analysis showed that Gp94 most likely functions as a monomer (**Error! Reference source not found.**B). Its predicted structure was compared to other protein structures using Foldseek^32^, but no similarity to proteins of known function was found, and therefore, no putative function could be proposed.

### Gp94 changes cell morphology through interference with peptidoglycan synthesis

To investigate the inhibitory effect of Gp94 over time, growth was followed using OD_600_ measurements and time-lapse microscopy, and compared with the control. The empty vector control grew exponentially for 10 to 12 hours, after which the stationary phase was reached, and no differences were observed between the induced and uninduced conditions (Figure 4A). ESBL098 transformed with pPS26::*gp94* seemed to grow less well than the empty vector control, even in the absence of the inducer. This could indicate some leakiness of the promoter, although no inhibition was seen when the strain was spotted on plates without inducer (Figure 2). When expressing Gp94, both OD_600_ measurements and time-lapse microscopy showed that cells were growing as the control for two to three hours after induction, after which growth stopped completely (Figure 4B and Video S1). This resulted in a smaller colony area of ESBL098 expressing *gp94* than the empty vector control (Figure 4C). As observed ealier, cells became bloated and also developed an uneven appearance, which could be due to cytoplasmic density differences (Figure 4D-E). Cells did not lyse visibly, but live-dead staining showed that cells began dying after five hours of Gp94 production, with a peak in cell death at approximately 19 hours (Figure 4C, Figure 4F, and Video S2). After eight hours of induction, the observed OD_600_ increased again, possibly due to colonies that had acquired resistance against Gp94. This was also observed during time-lapse microscopy, in which new colonies formed in one of three replicates, each originating from a single cell and resembling the control (Video S1). This confirms the occurrence of resistance in a limited number of cells. Five resistant colonies were further investigated, but a colony PCR revealed that resistance was caused by an approximately 1 kb insertion in the plasmid encoding *gp94* rather than a mutation of the bacterial target (**Error! Reference source not found.**). These results show that Gp94 production alters cell morphology and causes cell death.

**Figure 4:**
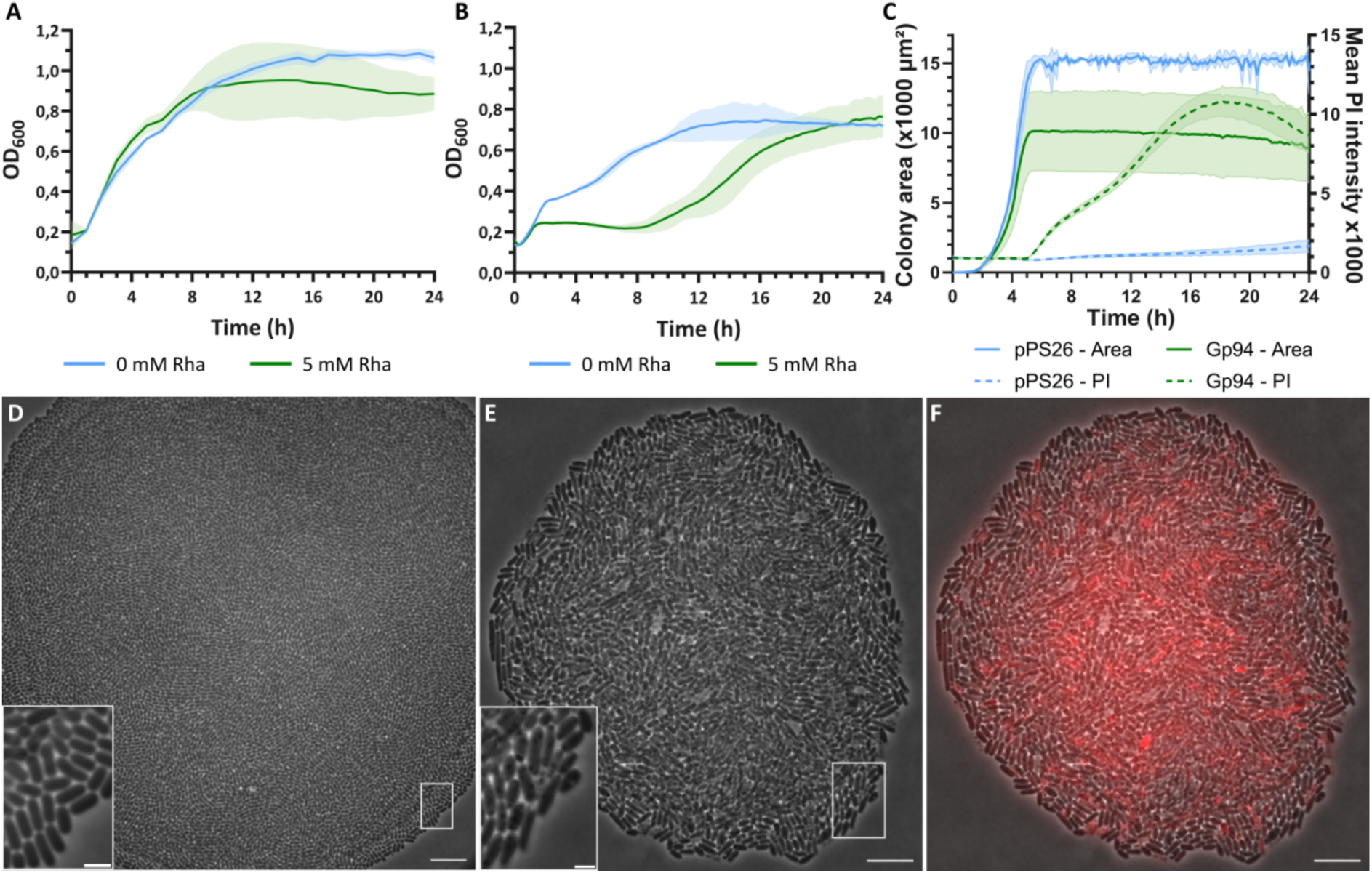
Expression of Gp94 completely inhibits growth of ESBL098. The growth of ESBL098 transformed with A) the empty vector pPS26 and B) pPS26::*gp94* was followed for 24 h in liquid medium without (blue) and with (green) rhamnose. The mean and standard deviation obtained from biological triplicates are shown. C) The detected colony area and the mean propidium iodide (PI) signal intensity during time-lapse microscopy of ESBL098 pPS26 and ESBL098 pPS26::*gp94* grown for 24 h in the presence of rhamnose. The mean and standard deviation obtained from triplicates are shown. D-E) Time-lapse microscopy of ESBL098 transformed with D) pPS26 or E) pPS26::*gp94* after 12 h of growth in the presence of rhamnose. F) Overlay of E with propidium iodide staining, showing dead cells in red. Scale bar = 10 µm. Scale bar insert = 2 µm. See also Video S2.

During time-lapse microscopy, it was observed that cells expressing Gp94 presented a distinct morphology. To gain a better understanding of the mechanisms underlying these morphological changes caused by Gp94 expression, the cell membrane, sites of active peptidoglycan synthesis, and the DNA were stained and visualized by fluorescence microscopy at different time points after induction. After one hour of Gp94 expression, no differences could be observed in any of the stained structures (**Error! Reference source not found.**A). However, after two hours of Gp94 expression, cells no longer formed septa, indicating arrested cell division (Figure 5A). Moreover, the location of peptidoglycan synthesis differed. Plotting the location of peptidoglycan synthesis foci onto an average cell for both the control and cells expressing Gp94 revealed differences in their spatial distribution (Figure 5B). In the empty vector control, foci were primarily located at the center of the cell, where the septum is usually formed. In cells expressing Gp94, foci were located at the periphery of the cell, and they were asymmetrically distributed along the length of the cell. This indicates that Gp94 expression interferes with the localization of peptidoglycan synthesis. After two hours of Gp94 expression, foci were also observed in the cell membrane, although this observation is likely related to cell death rather than a direct effect of Gp94. Similar observations, although more pronounced, were made after four hours of Gp94 expression (**Error! Reference source not found.**B). Differences in the DNA could not be observed at any point, apart from some variations in signal intensity between cells; however, it is unclear what causes these differences.

**Figure 5:**
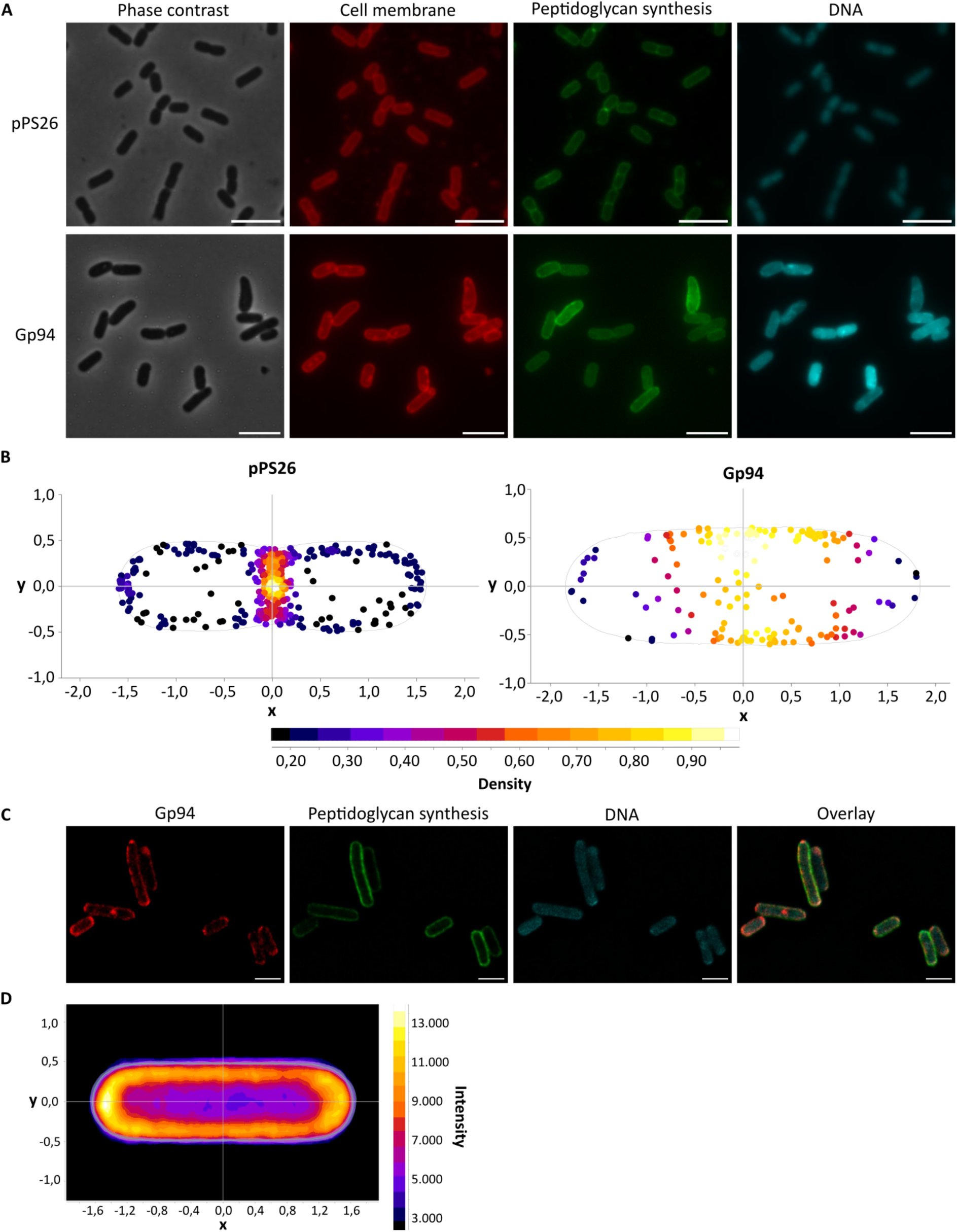
Expression of Gp94 induces changes in the location of peptidoglycan synthesis in ESBL098. A) Phase contrast and fluorescent images staining the cell membrane, peptidoglycan synthesis, and DNA of ESBL098 transformed with the empty vector or pPS26::*gp94* after two hours of induction. Representative images from 15 images for each condition are shown. Scale bar = 5 µm. B) The location of active peptidoglycan synthesis after two hours of induction was compared between the empty plasmid control (n = 627 maxima detected in 3452 cells) and Gp94 expression (n = 256 maxima detected in 2048 cells) in ESBL098. C) Fluorescent microscopy images after 4 h of Gp94 expression in ESBL098 showing the location of Gp94 in the cell, peptidoglycan synthesis, DNA, and an overlay of all three. Scale bar = 2 µm. D) Heatmap of the Gp94 signal intensity distribution in the cell (n = 183 cells).

To further investigate whether the changes in peptidoglycan synthesis are directly linked to Gp94 expression, the cellular location of Gp94 was determined through the fusion with a HaloTag, which can bind a fluorescent ligand. This revealed that Gp94 is localized to the cell periphery and overlays with the sites of peptidoglycan synthesis (Figure 5C-D). These findings support that Gp94 interferes with the localization of peptidoglycan synthesis.

### Growth inhibition by Gp94 can be complemented by the overexpression of MreBCD

Other phage proteins, such as the T7 protein Gp0.6 and the *E. coli* toxin YeeV, cause growth inhibition and morphological changes similar to those of Gp94 and have been shown to interact with MreB^22,33^. Therefore, we speculated that Gp94 interferes with a protein in the key elongasome complex, the MreBCD complex, a hypothesis supported by the localization of Gp94 at the cell periphery and its effect on peptidoglycan synthesis. To test this hypothesis, MreB, MreC, and MreD were overexpressed individually and in combination to investigate whether they could complement the effect of Gp94. Expression of MreB, MreD, and MreBCD did not inhibit growth, while MreC had a strong growth-inhibitory effect on when expressed individually (Figure 6A),. Gp94 expressed alone inhibited growth, as observed previously. However, when MreB, MreD, and MreBCD were expressed together with Gp94, more dense growth was observed compared to expression of Gp94 alone (Figure 6A), indicating that these genes can partially complement the growth-inhibitory effect of Gp94. Interestingly, the impact of MreB expressed individually or in combination with MreCD was similar. Overexpression of MreC in combination with Gp94 did not result in complementation, likely due to the toxic effects of MreC. Phase contrast microscopy confirmed that, upon expression of Gp94 and MreBCD, the cell morphology was partially restored to a rod-shaped morphology (Figure 6B). Overall, the complementation assays suggest that MreB, MreD, and the MreBCD complex partially complement the inhibitory effect of Gp94.

**Figure 6:**
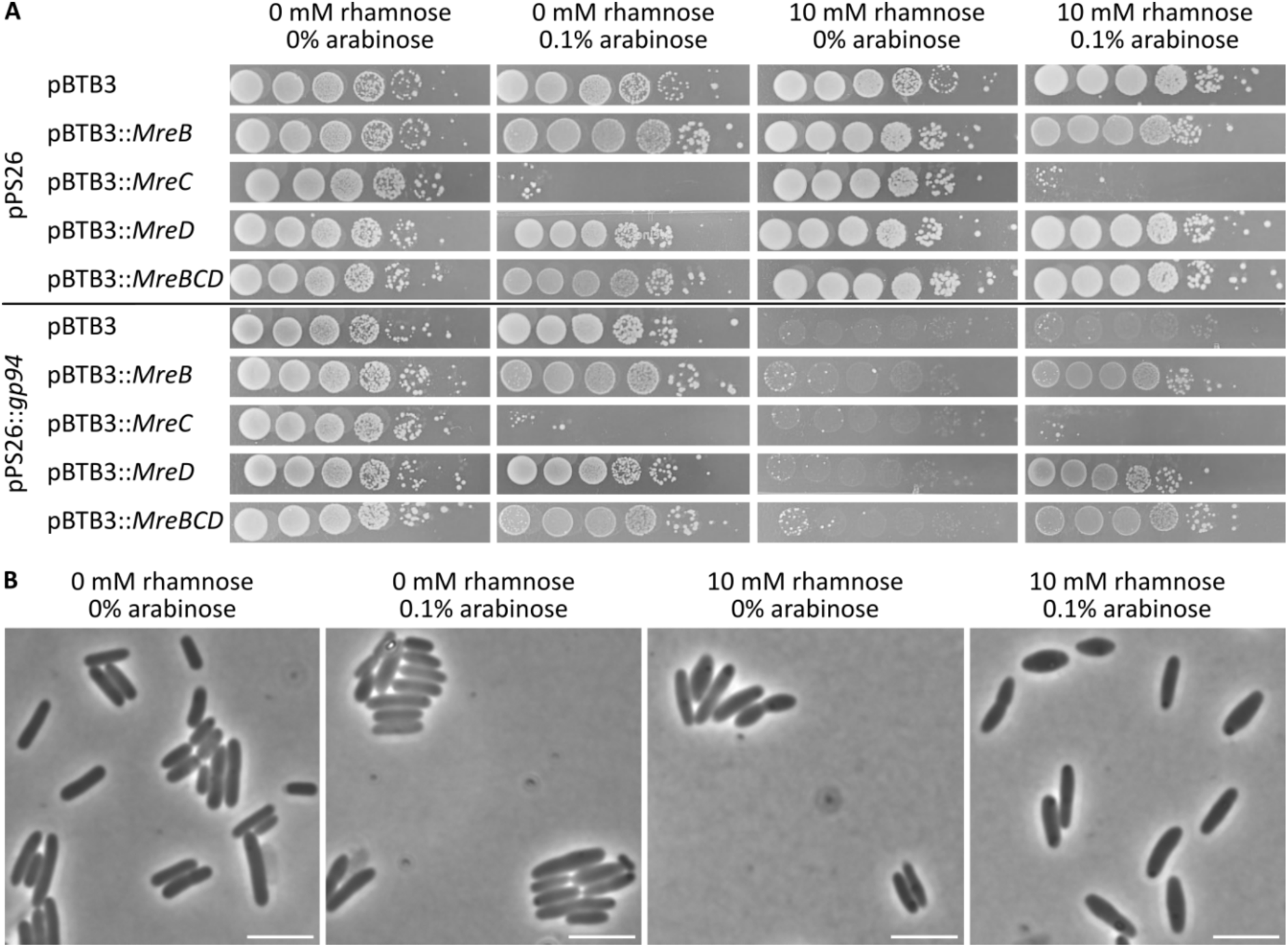
MreBCD complements the growth-inhibitory effect and the morphological changes caused by Gp94. A) Tenfold dilutions of Stellar^TM^ transformed with either the empty vector pPS26 or pPS26::*gp94* and pBTB-3 encoding the respective gene were spotted on LA plates with and without 10 mM rhamnose and 0.1% arabinose. B) Phase-contrast microscopy of Stellar^TM^ pPS26::*gp94* pBTB3::*MreBCD* after two hours of growth in the presence and absence of rhamnose and arabinose. Scale bar = 5 µm.

### Gp94 interacts with MreB, MreC, and MreD and is renamed Emp, elongasome-modulating peptide

The complementation assay showed that the growth inhibitory effect of Gp94 was partially rescued by expressing MreB, MreD, and MreBCD *in trans*. A bacterial two-hybrid assay was used to determine whether Gp94 directly interacts with MreBCD complex^34^. Together with the leucine zipper (ZIP) domain as a negative control, FtsZ was used to determine whether Gp94 interacts only with proteins in the elangosome and not with FtsZ, which is part of the divisome. The positive control pUT18C_ZIP and pTK25_ZIP yielded distinct blue colonies, whereas the negative control co-transformants (pUT18_Gp94 with pTK25_ZIP) remained white (Figure 7A). In line with our hypothesis, Gp94 showed no interaction with FtsZ, confirming that it does not nonspecifically bind to general cell-division factors. In contrast, Gp94 generated positive interaction signals with MreB, MreC and MreD, but was orientation specific. Only when the T25 fragment was fused to the C-terminus, an interaction with Gp94 was observed. The interaction with MreB produced a more intense blue color than the interactions with MreC and MreD. (Figure 7A). In conclusion, we propose that Gp94 perturbs elongasome-associated peptidoglycan synthesis in an MreBCD-dependent manner and rename it Emp (Elongasome-Modulating Peptide).

**Figure 7.**
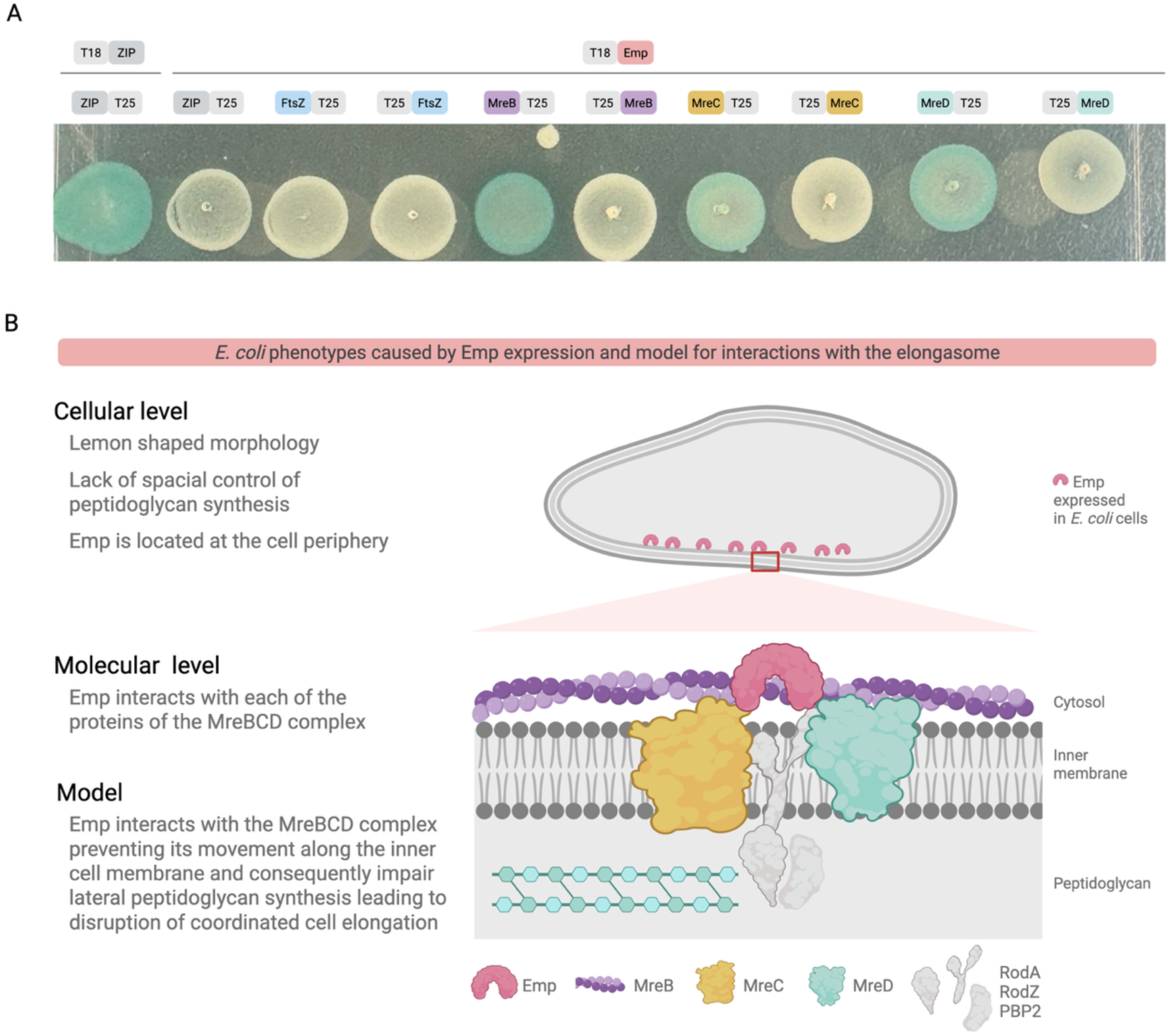
Emp interacts with elongation proteins MreB and MreD. **A)** *E. coli* BTH101 cells carrying Emp-T18 fusion constructs together with the FtsZ, MreB, MreC, and MreD T25 fusion partners were spotted onto X-gal plates to investigate protein-protein interaction. The leucine zipper pair (ZIP-T18/T25-ZIP) served as a positive control, and Emp-T18 together with T25-ZIP served as a negative control. A strong interaction between Emp and MreB was detected in an orientation-dependent manner. A weaker interaction was observed between Emp and MreC, and MreD. Emp did not interact with FtsZ in any fusion orientation. **B)** Proposed model for Emp-mediated disruption of elongasome function. When Emp is expressed in *E. coli*, cells exhibit a lemon-shaped morphology and lack specific control of peptidoglycan synthesis. Also, Emp is located at the cell periphery. At the molecular level, Emp interacts with MreBCD, which, under normal conditions, directs peptidoglycan synthesis along the lateral cell wall, maintaining rod-shaped morphology. Emp interacts with each of the MreBCD components, preventing MreB movement along the inner cell membrane and consequently impairing lateral peptidoglycan synthesis. This disrupts coordinated cell elongation and normal rod morphology, leading to lemon-shaped cells and eventual cell lysis.

Expression of Emp (Gp94) induced a characteristic morphological phenotype in *E. coli*, where cells progressively lost their normal rod-shaped morphology, adopted a lemon-shaped appearance, and ultimately lysed. Together with the fluorescent microscopy and bacterial two-hybrid assay results, these observations suggest that Emp interferes with the MreBCD elongasome complex, disrupting the spatial organization of peptidoglycan synthesis required to maintain rod shape. We therefore propose that Emp associates with elongasome components, leading to aberrant cell wall synthesis and subsequent cell envelope failure (Figure 7B).

## Discussion

During the early stage of phage infection, phages reprogram the metabolism of their host and direct it towards the production of new phage progeny. Despite growing knowledge of the phage proteins involved in this process, the functions of many early-expressed proteins remain unknown. This is especially true in less-studied phage genera such as *Mosigvirus*. In this study, we identified early-expressed proteins of *Mosigvirus* AV110 that are potentially involved in host reprogramming, based on their sequence similarity to the known early-expressed proteins of *Tequatrovirus* T4. This resulted in the identification of 63 putative early-expressed proteins. Of the 90 known early-expressed proteins of T4, 27 did not have a homolog in AV110. These proteins mainly encode for hypothetical proteins, but also two homing endonucleases, the ModA RNA polymerase ADP-ribosylase, the Arn inhibitor of MrcBC restriction, and the Stp activator of host PrrC lysyl-tRNA endonucleases. This indicates that these functions are either specific to *Tequatrovirus* members or carried out by a protein of AV110 unrelated at the sequence level.

Of the 63 proteins identified in AV110, 45 had an unknown or poorly characterized function. The effect of recombinant expression of these 45 proteins was tested against the ESBL/AmpC *E. coli* strain ESBL098, identifying six proteins, or 13% of the tested proteins, with a growth-inhibitory effect. This indicates that they interfere with important host processes for bacterial growth and might be involved in host reprogramming. The number of proteins with an antibacterial effect is slightly lower than in similar systematic screenings for antibacterial proteins from phages LUZ7^35^, T5^36^, and T7^22^, in which 30%, 23%, and 35%, respectively, exhibited a growth-inhibitory effect. This difference can be explained by excluding proteins with a known function from the screening, since this also included eight proteins known to have a growth-inhibitory effect. Together, this resulted in 22% of the identified early-expressed proteins with a growth-inhibitory effect. Nevertheless, AV110 may encode additional early-expressed proteins with antibacterial properties that require, for example, a chaperone for proper folding or formation of a complex to be active. For instance, the T4 DNA replication helicase (Gp41) requires interaction with the helicase loader protein (Gp59) to unwind the DNA and translocate along Gp32-coated ssDNA^37^. Due to the design of the screening assay, such proteins are likely to be missed. In addition, early-expressed proteins of *Mosigvirus* phages, which are not found in *Tequatrovirus* phages, are also missed. In this regard, it could be beneficial to perform RNA sequencing and mass spectrometry at different time points during the course of the infection to obtain a more comprehensive overview of the transcriptional and translational landscape of AV110, including the early-expressed proteins, as is avalible for T4^14^.

Of the 45 early-expressed proteins, six inhibited the growth of ESBL098. When tested across a genetically diverse panel of ESBL/AmpC *E. coli* strains, Gp214 and Gp263 only affected the growth of ESBL098, whereas Gp208 inhibited ESBL098 and ESBL049. Notably, Gp208 did not inhibit the growth of ESBL032, despite ESBL032 and ESBL049 belonging to the same sequence type (ST-23), for which a similar effect might have been expected. This suggests that these proteins interfere with a process unique to ESBL098 (and ESBL049). Alternatively, the targeted function could be performed by a variant of the ESBL098 protein or by a structurally distinct protein that does not interact with the phage protein. It is also possible that the growth-inhibitory effect is compensated for by an alternative protein or pathway.

The early-expressed protein Gp94 (Emp) was identified as the protein with the most substantial growth-inhibitory effect in diverse ESBL/AmpC *E. coli* strains. Upon recombinant expression, cell morphology was altered, resulting in bloated, lemon-shaped cells. A similar growth-inhibitory effect and morphological changes were observed after expressing gp0.6 from phage T7^22^. Using a mutant screening, MreB was identified as the target of this protein. MreB, as well as MreC and MreD, are key proteins of the elongasome, which is important for localized peptidoglycan synthesis during cell growth^38^. MreB forms large filaments that direct the insertion of new peptidoglycan molecules in the proper location, thereby maintaining the rod-shaped morphology of the cell. The membrane-spanning proteins MreC and MreD are recruited by MreB to bridge and regulate interactions between cytoplasmic MreB and extracellular penicillin-binding proteins PbP2 in the periplasm. Here, we found that the growth-inhibitory effect of Emp was partially complemented by the overexpression of MreB, MreD and the whole MreBCD complex. Additionally, the bacterial two-hybrid assay demonstrated that Emp interacts with MreB, MreC, and MreD, but suggests a stronger interaction with MreB than MreC and MreD. The *E. coli-*encoded toxin YeeV causes morphological changes similar to those observed with Emp^33^. A pull-down experiment and yeast two-hybrid assay showed that YeeV interacts with MreB and the cell-division protein FtsZ. However, while YeeV interacts with both MreB and FtsZ, Emp does not interact with FtsZ, further suggesting that Emp employs a different molecular mechanism than YeeV. The interaction of Emp with the MreBCD complex suggests that the protein targets the core of the elongasome rather than the divisome. The observed lemon-shaped morphology and subsequent lysis following Emp expression are therefore consistent with disruption of MreBCD-dependent control of peptidoglycan synthesis. We propose that Emp associates with the MreBCD complex and perturbs the spatial organization of cell wall assembly, resulting in aberrant peptidoglycan insertion, loss of rod-shaped growth, and ultimately cell wall failure. Still, despite the similar phenotypical effects of Emp, Gp0.6, and YeeV and their interaction with MreB, they do not share any sequence or structural similarity. This suggests that their molecular interactions interfere with elongation and cell division through distinct mechanisms. However, interacting with and inhibiting cell division and elongasome proteins could provide an energy advantage for the phage during infection. Additionally, it has been suggested that inhibiting MreB loosens the cell wall, which may facilitate the smoother release of new virions^22^. However, further studies are needed to confirm this.

Interference with the peptidoglycan layer is the mode of action of the commonly used β-lactam antibiotics. They covalently bind to the penicillin-binding proteins (PBPs), which are responsible for crosslinking peptidoglycan chains. By binding to the PBP, β-lactam antibiotics inhibit this crosslinking activity, resulting in a weakened cell wall and cell death^39^. In *E. coli*, β-lactam antibiotics can induce different phenotypes depending on the targeted PBP. Penicillin, which interferes with PBP2, typically causes the formation of spherical cells, while third-generation cephalosporins like moxalactam interfere with PBP3, resulting in cell filamentation^40^. However, Emp affected *E. coli* morphology differently, causing bloated cells and targeting MreB-dependent peptidoglycan synthesis. Therefore, identifying the molecular interactions of Emp and MreB could allow the development of novel antibacterials targeting MreB. Since MreB is essential for growth and is not found in eukaryotic cells, mimicking the effect of Emp with a small molecule, as exemplified by proteins with a growth-inhibitory impact of *S. aureus* phages^13^, could enable the development of a novel antibacterial agent, potentially effective against bacteria resistant to β-lactam antibiotics.

In conclusion, these findings highlight the potential of early-expressed phage proteins with antibacterial properties to elucidate interactions between phages and their hosts, which could be exploited in future discovery new antibacterial targets.

## Supporting information

All supplementary information

## Methods

### Key resources table

| REAGENT or RESOURCE | SOURCE | IDENTIFIER |
| --- | --- | --- |
| Bacterial and virus strains |  |  |
| ESBL032 | Vitt et al. 2023<sup>2</sup> | SAMEA3993582 |
| ESBL049 | Vitt et al. 2023<sup>2</sup> | SAMEA3993608 |
| ESBL054 | Vitt et al. 2023<sup>2</sup> | SAMEA3993615 |
| ESBL080 | Vitt et al. 2023<sup>2</sup> | SAMEA4058391 |
| ESBL087 | Vitt et al. 2023<sup>2</sup> | SAMEA4058402 |
| ESBL098 | Vitt et al. 2023<sup>2</sup> | SAMEA4058413 |
| ESBL115 | Vitt et al. 2023<sup>2</sup> | SAMEA4058431 |
| ESBL154 | Vitt et al. 2023<sup>2</sup> | SAMEA104206015 |
| BTH101 | EUROMEDEX | EUK001 |
| Stellar™ Competent Cells | Takara Bio | Cat#636766 |
| Mosigvirus AV110 | Vitt et al. 2024<sup>24</sup> | GenBank: OR352940.1 |
| Chemicals, peptides, and recombinant proteins |  |  |
| L-arabinose | Thermo Fisher Scientific | Cat#365180250 |
| L-rhamnose | Millipore | Cat#83650 |
| DAPI | Invitrogen | Cat#D1306 |
| Janelia Fluor® 646 HaloTag® Ligand | Promega | Cat#HT1060 |
| Nile-Red | Invitrogen | Cat#N1142 |
| OGDA | Tocris Bioscience | Cat#7408 |
| Propidium iodide | Invitrogen | Cat#P1304MP |
| Critical commercial assays |  |  |
| CloneAmp HiFi PCR Premix | Takara Bio | Cat#639298 |
| DreamTaq Green PCR Master Mix | Thermo Fisher Scientific | Cat#K1082 |
| GeneJET PCR Purification Kit | Thermo Fisher Scientific | Cat#K0702 |
| GeneJET Plasmid Miniprep Kit | Thermo Fisher Scientific | Cat#K0502 |
| InFusion Snap Assembly Master Mix | Takara Bio | Cat#638948 |
| Bacth kit | EUROMEDEX | EUK001 |
| Oligonucleotides |  |  |
| Primers for cloning | See **Error! Reference source not found.** | N/A |
| Recombinant DNA |  |  |
| pPS26 | Calero et al. 2016<sup>41</sup> | N/A |
| pPS26-gp0033 – pPS26-gp0264 | This study | N/A |
| pPS26-gp94-Halotag | This study | N/A |
| pPS26-alc | This study | N/A |
| pBTB3 | Lynch et al. 2006<sup>42</sup> | Addgene #22819 |
| pBTB3-mreB | This study | N/A |
| pBTB3-mreC | This study | N/A |
| pBTB3-mreD | This study | N/A |
| pBTB3-mreBCD | This study | N/A |
| pUT18-gp94 | This study | N/A |
| pUT18C-ZIP | EUROMEDEX | EUK001 |
| pTK25-ZIP | EUROMEDEX | EUK001 |
| pTK25-FtsZ | This study | N/A |
| pNTK25-FtsZ | This study | N/A |
| pTK25-MreB | This study | N/A |
| pNTK25-MreB | This study | N/A |
| pTK25-MreC | This study | N/A |
| pNTK25-MreC | This study | N/A |
| pTK25-MreD | This study | N/A |
| pNTK25-MreD | This study | N/A |
| Software and algorithms |  |  |
| CLC Workbench 23 | Qiagen | N/A |
| Fiji | Schindelin et al. 2012<sup>43</sup> | N/A |
| LoVis4u | Egorov et al. 2025<sup>29</sup> | N/A |
| MicrobeJ | Ducret et al. 2016<sup>44</sup> | N/A |

### Experimental model and study participant details

#### Bacterial strains and growth conditions

The bacterial strains, phages, and vectors used in this study are indicated in the Key resources table. All *E. coli* strains were streaked out on LA plates, when necessary supplemented with 50 µg/ml kanamycin and/or 25 µg/ml chloramphenicol, and incubated overnight at 37°C. Next, a single colony was picked, inoculated in LB, when necessary supplemented with 50 µg/ml kanamycin and/or 25 µg/ml chloramphenicol and incubated overnight at 37°C with shaking at 180 rpm, unless otherwise specified.

### Method details

#### Identification of early-expressed proteins

The early-expressed proteins of T4 were identified based on transcriptome and proteome data^14^. All genes for which the transcript was detected early and the protein was detected early or NA were selected, resulting in 90 genes. Next, the amino acid sequences of the 90 early-expressed T4 genes were extracted and aligned to the amino acid sequences of all CDS of AV110 (OR352940.1) in CLC Main Workbench 23 (Qiagen). Proteins with sufficient sequence identity or similarity, a limited number of gaps, and a similar length were identified as an early-expressed protein of AV110.

#### Cloning of early genes

The DNA of phage AV110 was extracted by treating a filtered phage stock with 10 µg/ml RNase and 20 µg/ml DNase for 1 h at 37°C, followed by the addition of 20 mM EDTA and a proteinase K (50 µg/ml) treatment for 2 h at 56°C. Next, the DNA was purified using the Genomic DNA Clean & Concentrator-10 Kit (ZymoResearch) according to the manufacturer’s instructions. Primers were designed for all identified early-expressed genes with a 15 bp overhang complementary to the insertion site in the vector pPS26 (**Error! Reference source not found.**). Each gene was amplified and the vector pPS26 was linearized with the addition of a ribosome-binding site with PCR using the CloneAmp HiFi PCR Premix (Takara). Next, the size of the PCR product was verified on a 1% agarose gel and the PCR product was purified using the GeneJET PCR Purification Kit (Thermo Scientific). The purified PCR product of each early-expressed gene was mixed with the linearized vector in a 200 ng:100 ng ratio, together with the InFusion Snap Assembly Master Mix (Takara). The reaction mixture was incubated at 50°C for 15 min. Next, Stellar^TM^ Competent Cells (Takara) were transformed with the mixture by heat-shock at 42°C for 1 min, followed by the addition of 500 µl SOC medium and incubation at 37°C with shaking (180 rpm). Next, dilutions of the transformation mixture were spread on LA plates with kanamycin (50 µg/ml) and incubated overnight at 37°C. Multiple colonies of each gene were picked, and the insert size was verified by colony PCR using primers complementary to the backbone of the vector flanking the insert (**Error! Reference source not found.**) and the DreamTaq Green PCR Master Mix (Thermo Scientific). The size of the PCR products was verified on a 1% agarose gel. Plasmids with the correct insert size were purified from an overnight culture using the GeneJET Plasmid Miniprep Kit (Thermo Scientific) according to the manufacturer’s instructions and the sequence of each plasmid was verified by Plasmidsaurus using nanopore sequencing. Next, electrocompetent ESBL/AmpC *E. coli* strains were transformed with 30 ng of the plasmid via electroporation using the following settings: 1.8 kV, 25 µF, and 200 Ω, followed by the addition of 1 ml LB and incubation at 37°C with shaking (180 rpm) for 1 h. Cells were spread on LA plates with kanamycin and incubated overnight at 37°C. A single colony was picked and the insert size was again verified using colony PCR as before.

#### Growth-inhibition assay

The growth-inhibitory effect of each early-expressed gene was tested in ESBL098 in triplicate. Starting from an overnight culture of the ESBL098 strain encoding each gene induvidually, a tenfold dilution series was made in LB with kanamycin and 10 µl of the undiluted culture and each dilution until 10^-5^ were spotted on LA plates with kanamycin, with or without 5 mM rhamnose. The plates were incubated overnight at 37°C, after which the growth-inhibitory effect was assessed.

#### Growth curves

A growth curve of ESBL098 transformed with the empty vector pPS26 or pPS26::*gp94* was constructed in triplicate by incubating a single colony of each strain in LB with kanamycin to an OD_600_ of 0.2. Next, a twofold dilution of the culture was made in a 96-well plate in LB with kanamycin with or without rhamnose at a final concentration of 5 mM. Growth curves were obtained by incubating the plate in a PowerWave XS microplate reader (BioTek) at 37°C, with OD_600_ measurements every 15 min for 24 h.

#### Phase contrast and fluorescent microscopy

Phase contrast microscopy was performed using ESBL098 transformed with each growth-inhibitory gene after four hours of incubation in the presence of rhamnose. For ESBL098 pPS26::*gp94*, additional phase contrast and fluorescence microscopy images were obtained after one and two hours. An overnight culture of each strain was diluted to an OD_600_ of 0.05 and incubated for 30 min at 37°C with shaking (120 rpm). Next, 5 mM rhamnose and 10 µg/ml Nile Red were added, after which the culture was incubated for 1, 2, or 4 hours. Next, 100 µM OGDA and 5 µg/ml DAPI were added and the culture was incubated for 3 min and pelleted for 3 min at 4000 rpm. The pellet was resuspended in 4.6% formaldehyde and stored overnight at 4°C. The next day, the fixed cells were pelleted (4000 rpm, 3 min) and resuspended in PBS. A 1.2 % agarose pad in PBS was made on which 1.5 µl of the fixed cells was spotted. The cells were covered with a coverslip and imaged using an inverted Olympus IX83 microscope with a 1.4 N.A. 100x oil-immersion objective. Fifteen phase-contrast and fluorescent images were acquired for each strain and time point using the U-FMCHE (Nile Red), U-FBNA (OGDA), and U-FUNA (DAPI) filter cubes. The images were processed and analyzed in Fiji^43^ using the ImageJ plug-in MicrobeJ^44^.

#### Time-lapse microscopy and live-dead staining

Time-lapse microscopy was performed with and without live-dead staining with ESBL098 transformed with the empty plasmid pPS26 and pPS26::*gp94*. An overnight culture was diluted 1/100 and incubated for 1 h at 37°C. Next, a 1.2 % agarose pad in LB with kanamycin, 5 mM rhamnose, and, when necessary, 1 µM propidium iodide was prepared in a 1.7×2.8 cm Gene Frame (Thermo Scientific) in which eight wells were created using another Gene Frame. The culture was diluted to an OD_600_ of 0.01 and 1 µl of the diluted bacterial culture was spotted on the agarose pad and dried before a cover slip was applied. An inverted Olympus IX83 microscope with a 1.4 N.A. 100x oil-immersion objective was used with a cellVivo incubator module set at 37°C. Three colonies were selected for each strain. Phase contrast images were acquired every 5 min for 25 h with 20 ms exposure time and a sCMOS Photometric Prime camera. When propidium iodide was added, phase contrast images were acquired every 10 min for 27 h, together with a fluorescent image. The images were processed in Fiji^43^.

#### Protein localization

Overnight cultures of ESBL098 transformed with the empty vector pPS26 and pPS26::*gp94* fused at the C-terminal end with a HaloTag were diluted to an OD_600_ ∼ 0.001 in LB with kanamycin and were grown at 37°C for one hour, before adding 5 mM rhamnose for induction of *gp94*. Cells were induced for a total of four hours, where cells were stained with 100 µM OGDA (ToCris Biotechne) and 1 µM HaloTag ligand JF646 (Promega) one hour prior to fixation. Cells were stained with 1 µg/mL DAPI five minutes prior to fixation. After four hours of incubation, cells were briefly pelleted, washed in PBS, and fixed with 4% PFA for 20 minutes at room temperature. Cells were washed in PBS, resuspended in PBS, and spotted on 1.2 % PBS agarose pads with 1:10 ProLong Glass anti-fade mountant media (Thermo Fischer Scientific) and sealed with a coverslip. Cells were imaged on an Abberior STED microscope (Abberior Instruments), with a 60x 1.4 N.A. oil immersion objective, using a matrix detector for the green 488 channel. A STED laser was applied to the red 647 channel with 15 nm pixel size, while the blue 405 channel was imaged using standard confocal imaging. Images were appropriately deconvolved for intensity (red and blue channels), while the green channel was reconstructed using matrix deconvolution. Images were further analyzed in Fiji^43^ and with the ImageJ plug-in MicrobeJ^44^.

#### Complementation assay

Electrocompetent Stellar^TM^ pPS26::*gp94* was transformed with pBTB3::*MreB*, pBTB3::*MreC*, pBTB3::*MreD*, and pBTB3::*MreBCD* and incubated overnight at 37°C on LA plates with 50 µg/ml kanamycin and 25 µg/ml chloramphenicol, after which a single colony of each strain was picked. A growth-inhibition assay was performed as earlier using the following conditions: no inducer, 10 mM rhamnose, 0.1% arabinose, and 10 mM rhamnose and 0.1% arabinose. The effect on cell morphology was assessed by growing the cultures for 2 h at 37°C with shaking in LB with 50 µg/ml kanamycin and 25 µg/ml chloramphenicol without inducer, 10 mM rhamnose, 0.1% arabinose, and 10 mM rhamnose and 0.1% arabinose. Next, 1 µl was spotted on an agarose pad and sealed with a coverslip. Phase contrast images were acquired using a 1.3 N.A. 100x oil-immersion objective on an Axioplan 2 microscope (Zeiss).

#### Bacterial two-hybrid assay

To construct the bacterial two-hybrid vectors, the BACTH vectors, Gp94 and the target genes MreB, MreD, and FtsZ were amplified using specific primers (Table S3). The resulting PCR fragments were cloned into the BACTH vectors to generate pUT18_Gp94, pTK25_MreB, pTK25_MreC, pTK25_MreD, pTK25_FtsZ, pNTK25_MreB, pNTK25_MreC, pNTK25_MreD, and pNTK25_FtsZ using the In-Fusion Snap Assembly Master Mix as previously described. However, to suppress leaky basal transcription during plasmid propagation and cloning, all growth media were supplemented with 1% (w/v) D-glucose. For protein-protein interaction screens, electrocompetent *Escherichia coli* BTH101 cells were co-transformed with the corresponding pairs of T18- and T25-derived plasmids using electroporation as previously described. Co-transformants harboring pUT18_Gp94 combined with pTK25_ZIP served as negative controls, whereas cells co-transformed with pUT18_ZIP and pTK25_ZIP served as the positive control. To assay protein-protein interaction, single colonies of freshly transformed BTH101 cells were inoculated into 3 mL of LB medium (supplemented with appropriate antibiotics and 1% glucose) and grown for 5 h at 30°C with shaking at 180 rpm. A 10 μL aliquot of each culture was spotted onto LB agar plates containing 100 μg/mL ampicillin, 50 μg/mL kanamycin, 0.5 mM IPTG, and 40 μg/mL X-Gal. Plates were incubated overnight at 30°C and subsequently inspected for blue colony development, indicating functional interaction between the fused proteins.

### Quantification and statistical analysis

#### Image analysis

To prepare density distributions of the location of peptidoglycan synthesis in the cell, the ImageJ plug-in MicrobeJ 5.13p (1)^44^ was used. Parameters were optimized to detect individual bacterial cells in the phase-contrast images, after which foci in the cells were detected. To detect the foci of peptidoglycan synthesis, maxima were detected in the fluorescent microscopy images and parameters were optimized to reduce the background signal. The detected bacterial cells and maxima were manually verified. Next, an XYCellDensity plot was made in MicrobeJ. The number of analyzed cells and foci is indicated in the figure legend.

To prepare a heatmap of Gp94 expression in the cell, individual cells were detected using the ImageJ plug-in MicrobeJ 5.13p (1)^44^ and verified manually. A heatmap was constructed using the Shape Plot function and the fluorescent signal intensity corresponding to Gp94 expression.

## Acknowledgements

This work was supported by the Independent Research Fund Denmark (2035-00112B). RL is supported by the KU Leuven Project C1 ACES [C16/20/001].

## Author contributions

S.S.F. Van Overfelt: Conceptualization, Formal analysis, Investigation, Methodology, Visualization, Writing –original draft, Writing –review & editing A. N. Sørensen: Conceptualization, Investigation, Methodology, Supervision, Writing –review & editing

1. V. H. Mebus: Investigation, Methodology
2. J. R. Tornby: Formal analysis
3. J. Brewer: Conceptualization, Writing –review & editing
4. R. Lavigne: Conceptualization, Funding acquisition, Writing –review & editing
5. L. Brøndsted: Conceptualization, Funding acquisition, Project administration, Supervision, Writing –review & editing

## Declaration of generative AI and AI-assisted technologies

During the preparation of this work, the authors used Copilot in order to improve the readability. After using this tool, the authors reviewed and edited the content as needed and take full responsibility for the content of the publication.

