## Supplementary material for "The *Mosigvirus* AV110 Elongasome-Modulating Peptide (Emp) inhibits *Escherichia coli* growth by targeting Mre-dependent peptidoglycan synthesis": All supplementary information

**Table S1: Overview of the early proteins identified in AV110 and their T4 homolog, related to** Error! Reference source not found.**.** Early-expressed proteins in AV110 were identified by aligning the CDS of the T4 early proteins, identified by Wolfram-Schauerte *et al.* (2022)[14], to all AV110 CDS. Genes with a known function or growth-inhibitory effect on *E. coli* were excluded from the analysis.

| Locus tag | Product | T4 homolog | T4 function | T4 reference |
|---|---|---|---|---|
| AV110_CDS_0033 | Hypothetical protein | alt.-3 | alt.-3 conserved hypothetical protein | N/A |
| AV110_CDS_0039[1] | Virion structural protein | 30.3' | gp30.3' hypothetical protein | N/A |
| AV110_CDS_0040 | Hypothetical protein | 30.4 | gp30.4 conserved hypothetical protein | N/A |
| AV110_CDS_0041 | Hypothetical protein | 30.5 | gp30.5 hypothetical protein | N/A |
| AV110_CDS_0043 | Hypothetical protein | 30.7 | gp30.7 conserved hypothetical protein | N/A |
| AV110_CDS_0044 | Hypothetical protein | 30.8 | gp30.8 conserved hypothetical protein | N/A |
| AV110_CDS_0045 | Hypothetical protein | 30.9 | gp30.9 conserved hypothetical protein | N/A |
| AV110_CDS_0053 | Hypothetical protein | cd.2 | cd.2 conserved hypothetical protein | N/A |
| AV110_CDS_0060 | Hypothetical protein | pseT.1 | pseT.1 conserved hypothetical protein | N/A |
| AV110_CDS_0061 | Rz-like spanin | pseT.2 | rz-like spanin | N/A |
| AV110_CDS_0063 | Inhibitor of host transcription | alc | inhibitor of host transcription | [19,45] |
| AV110_CDS_0072 | Dihydrofolate reductase | frd | dihydrofolate reductase | N/A |
| AV110_CDS_0074 | Hypothetical protein | frd.2 | hypothetical protein | N/A |
| AV110_CDS_0077 | DNA helicase loader | 59 | DNA helicase loader | [46] |
| AV110_CDS_0087 | Holin | t | holin | [47] |
| AV110_CDS_0088 | Anti-sigma factor | asiA | anti-sigma factor | [48] |
| AV110_CDS_0092 | Hypothetical protein | arn.2 | hypothetical protein | N/A |
| AV110_CDS_0093 | Hypothetical protein | arn.3 | arn.3 conserved hypothetical protein | N/A |
| AV110_CDS_0094 | Hypothetical protein | arn.4 | arn.4 conserved hypothetical protein | N/A |
| AV110_CDS_0095 | MotA-like activator of middle period transcription | motA | motA-like activator of middle period transcription | [49,50] |
| AV110_CDS_0100 | Acridine resistance protein | ac | ac acridine resistance protein | N/A |
| AV110_CDS_0101 | Ndd-like nucleoid disruption protein | ndd | ndd-like nucleoid disruption protein | [51] |
| AV110_CDS_0102 | Hypothetical protein | ndd.1 | hypothetical protein | N/A |
| AV110_CDS_0108 | RIIA lysis inhibitor | rIIA | rIIA lysis inhibitor | [52,53] |
| AV110_CDS_0109 | Hypothetical protein | rIIA.1 | rIIA.1 hypothetical protein | N/A |
| AV110_CDS_0113 | FmdB-like transcriptional regulator | 39.2 | FmdB-like transcriptional regulator | [54] |
| AV110_CDS_0114 | goF mRNA metabolism modulator | goF | hypothetical protein | N/A |

| Locus tag | Product | T4 homolog | T4 function | T4 reference |
|---|---|---|---|---|
| AV110_CDS_0115 | Cef modifier of suppressor tRNAs | cef | cef modifier of supressor tRNAs | [55] |
| AV110_CDS_0118[1] | MotB-like transcriptional regulator | motB | motB-like transcriptional regulator | [56,57] |
| AV110_CDS_0122 | Homing endonuclease | dda.1 | dda.1 hypothetical protein | N/A |
| AV110_CDS_0123 | Srd anti-sigma factor | srd | srd anti-sigma factor | [58] |
| AV110_CDS_0124 | RNA polymerase ADP-ribosylase | modB | RNA polymerase ADP-ribosylase | [59] |
| AV110_CDS_0135 | Nucleoside triphosphate pyrophosphohydrolase | 56 | dUTPase | N/A |
| AV110_CDS_0140 | Hypothetical protein | 61.2 | gp61.2 hypothetical protein | N/A |
| AV110_CDS_0141 | Spackle periplasmic | sp | spackle periplasmic | [60] |
| AV110_CDS_0143 | Hypothetical protein | imm.1 | iduronate sulfatase | N/A |
| AV110_CDS_0144 | Dmd discriminator of mRNA degradation | dmd | dmd discriminator of mRNA degradation | [61] |
| AV110_CDS_0168 | Hypothetical protein | 46.1 | DUF5487 family protein | N/A |
| AV110_CDS_0177 | Hypothetical protein | 55.1 | gp55.1 hypothetical protein | [62] |
| AV110_CDS_0178 | Hypothetical protein | 55.2 | gp55.2 hypothetical protein | [63] |
| AV110_CDS_0180 | Hypothetical protein | 55.3 | gp55.3 hypothetical protein | N/A |
| AV110_CDS_0181 | Hypothetical protein | 55.4 | gp55.4 conserved hypothetical protein | N/A |
| AV110_CDS_0182 | Hypothetical protein | 55.5 | gp55.5 conserved protein of unknown function | N/A |
| AV110_CDS_0184 | Hypothetical protein | 55.6 | gp55.6 conserved hypothetical protein | N/A |
| AV110_CDS_0188 | Hypothetical protein | 55.8 | gp55.8 conserved hypothetical predicted membrane protein | N/A |
| AV110_CDS_0192 | Endonuclease VII | 49 | endonuclease VII | [64] |
| AV110_CDS_0199 | Hypothetical protein | nrdC.3 | hypothetical protein | N/A |
| AV110_CDS_0201 | Hypothetical protein | nrdC.8 | hypothetical protein | N/A |
| AV110_CDS_0202[1] | Internal virion protein | nrdC.9 | nrdC.9 conserved hypothetical protein | N/A |
| AV110_CDS_0204 | Hypothetical protein | nrdC.10 | hypothetical protein | N/A |
| AV110_CDS_0208 | Hypothetical protein | mobD.1 | mobD.1 conserved hypothetical protein | N/A |
| AV110_CDS_0214 | Hypothetical protein | mobD.2a | mobD.2a hypothetical protein | N/A |
| AV110_CDS_0224 | Endoribonuclease | regB | endoribonuclease | [65] |
| AV110_CDS_0229 | Hypothetical protein | vs.7 | hypothetical protein | N/A |
| AV110_CDS_0230[1] | Hypothetical protein | vs.8 | hypothetical protein | N/A |
| AV110_CDS_0236 | Endonuclease V N-glycosylase UV repair enzyme | denV | endonuclease V N-glycosylase UV repair enzyme | [66] |
| AV110_CDS_0250[1] | Hypothetical protein | e.4 | hypothetical protein | N/A |
| AV110_CDS_0251 | Hypothetical protein | e.5 | e.5 conserved hypothetical protein | N/A |
| AV110_CDS_0253 | Hypothetical protein | frd.3 | hypothetical protein | N/A |
| AV110_CDS_0255 | Hypothetical protein | e.8 | e.8 conserved hypothetical protein | N/A |
| AV110_CDS_0262 | Hypothetical protein | trna.2 | trna.2 conserved hypothetical protein | N/A |

| Locus tag | Product | T4 homolog | T4 function | T4 reference |
|---|---|---|---|---|
| AV110_CDS_0263 | Hypothetical protein | trna.3 | trna.3 conserved hypothetical protein | N/A |
| AV110_CDS_0264 | Hypothetical protein | trna.4 | hypothetical protein | N/A |

[1]The start and/or end position of these genes was altered in the latest annotation of the AV110 genome compared to the annotation that was used to predict early-expressed proteins and their growth-inhibitory effect (OR352940.1).

**Table S2: Effect of early phage proteins on the growth of ESBL098, related to** Error! Reference source not found.**.** Tenfold dilutions of ESBL098 transformed with the indicated phage gene encoded on the vector pPS26 were spotted on LA plates without and with rhamnose. Genes with a growth-inhibitory effect are highlighted in red. The empty plasmid pPS26 and the T4 gene *alc* are used as the negative and positive controls.

| **Locus tag** | **0 mM Rha** | **5 mM Rha** |
|---|---|---|
| AV110_CDS_0033 | | |
| AV110_CDS_0039 | | |
| AV110_CDS_0040 | | |
| AV110_CDS_0041 | | |
| AV110_CDS_0043 | | |
| AV110_CDS_0044 | | |
| AV110_CDS_0045 | | |
| AV110_CDS_0053 | | |
| AV110_CDS_0060 | | |
| AV110_CDS_0061 | | |
| AV110_CDS_0072 | | |
| AV110_CDS_0074 | | |
| AV110_CDS_0092 | | |
| AV110_CDS_0093 | | |
| AV110_CDS_0094 | | |
| AV110_CDS_0100 | | |
| AV110_CDS_0102 | | |
| AV110_CDS_0109 | | |
| AV110_CDS_0114 | | |
| AV110_CDS_0122 | | |
| AV110_CDS_0135 | | |
| AV110_CDS_0140 | | |
| AV110_CDS_0143 | | |
| AV110_CDS_0168 | | |
| AV110_CDS_0178 | | |
| AV110_CDS_0180 | | |
| AV110_CDS_0181 | | |
| AV110_CDS_0182 | | |
| AV110_CDS_0184 | | |
| AV110_CDS_0188 | | |

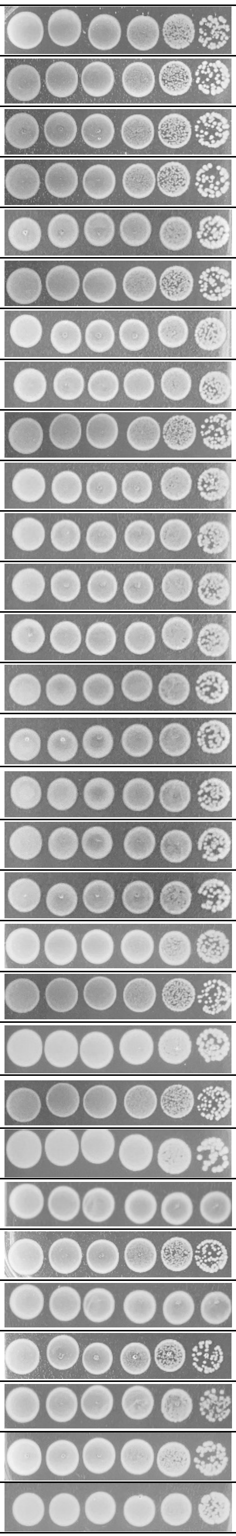

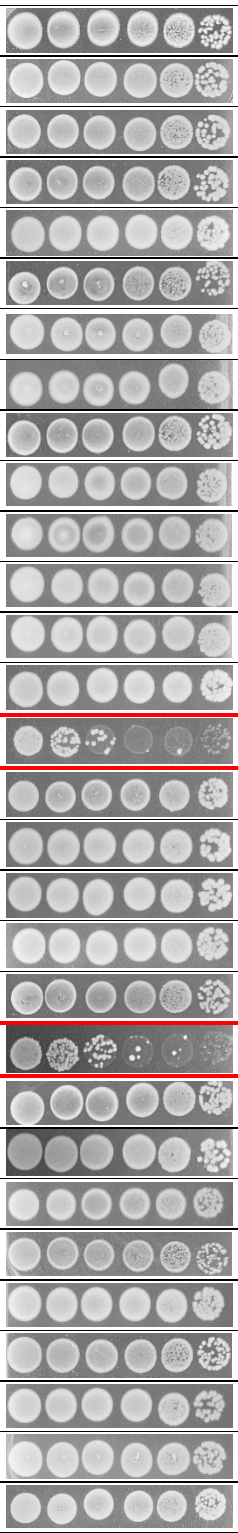

| **Locus tag** | **0 mM Rha** | **5 mM Rha** |
|---|---|---|
| AV110_CDS_0199 | | |
| AV110_CDS_0201 | | |
| AV110_CDS_0202 | | |
| AV110_CDS_0204 | | |
| AV110_CDS_0208 | | |
| AV110_CDS_0214 | | |
| AV110_CDS_0229 | | |
| AV110_CDS_0230 | | |
| AV110_CDS_0250 | | |
| AV110_CDS_0251 | | |
| AV110_CDS_0253 | | |
| AV110_CDS_0255 | | |
| AV110_CDS_0262 | | |
| AV110_CDS_0263 | | |
| AV110_CDS_0264 | | |
| AV110_CDS_0063[1] | | |
| AV110_CDS_0101[1] | | |
| AV110_CDS_0224[1] | | |
| pPS26 (neg. control) | | |
| T4 *alc* (pos. control) | | |

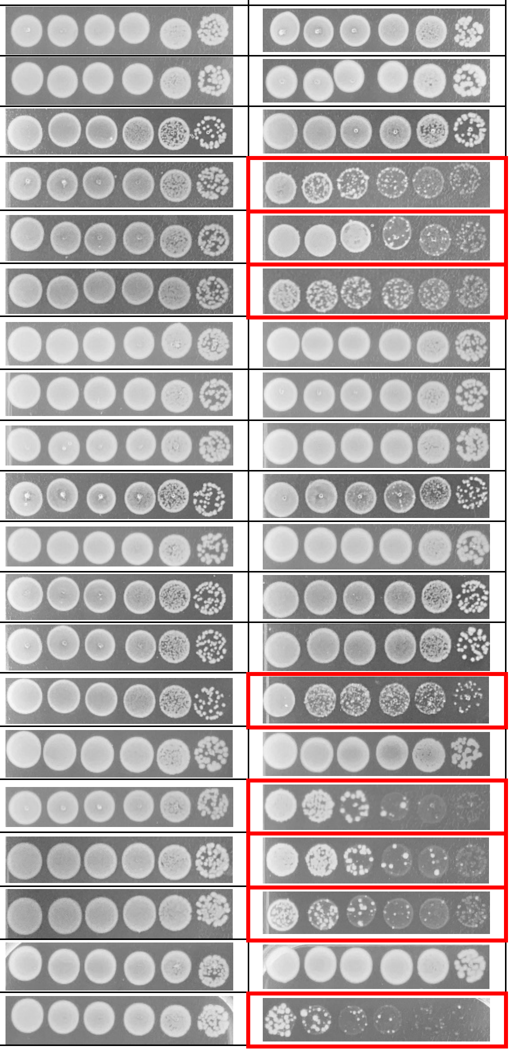

[1] These genes are homologs of T4 genes with a known growth inhibitory effect and were included as a proof-of-concept.

**Table S3: Overview of the primers used in this study, related to Methods.**

| Primer ID | Sequence (5' -> 3') |
|---|---|
| pPS26-lin_Fw | CACCTCAGCGTCGTGACTG |
| pPS26-lin_Rv | GGTATATTCCTCCTGCCTCAGCGAATTTCATTACGACC |
| pPS26_Fw | TAGTAATCACGAGGTCAGGT |
| pPS26_Rv | AGATGGAGTTCTGAGGTCA |
| T4-alc_Fw | GAGGAGGAATATACCATGGATTTACAACTTATTACTACTG |
| T4-alc_Rv | CACGACGCTGAGGTGTTACATGCATAAAGTTTTAATAACC |
| AV110_33-Fw | CAGGAGGAATATACCATGAAATCTATGCTTCGCTTTAATG |
| AV110_33-Rv | CACGACGCTGAGGTGTTATTTACGGAATGAAAGGAATGCT |
| AV110_39-Fw | CAGGAGGAATATACCGTGAAAAATGTTGAACAAC |
| AV110_39-Rv | CACGACGCTGAGGTGTCATAAAGAGTCTCTTAATAGG |
| AV110_40-Fw | CAGGAGGAATATACCATGTTGAATAAATTAATCCAGA |
| AV110_40-Rv | CACGACGCTGAGGTGTCAGACATCTTTAACACT |
| AV110_41-Fw | CAGGAGGAATATACCATGAAATTTTTAATAGCACAAACAG |
| AV110_41-Rv | CACGACGCTGAGGTGTTATTCAACATGTTTCAACCC |
| AV110_43-Fw | CAGGAGGAATATACCATGAACTACACCAACTTCGAAC |
| AV110_43-Rv | CACGACGCTGAGGTGTTAGATATCAAATTCGTTAAGAACT |
| AV110_44-Fw | CAGGAGGAATATACCATGAAAATCAATCTGAACTCATACA |
| AV110_44-Rv | CACGACGCTGAGGTGTTATGCAACTTTAATCCAATTGTCT |
| AV110_45-Fw | CAGGAGGAATATACCATGGCAAAACAAGCTAAAGCAAAG |
| AV110_45-Rv | CACGACGCTGAGGTGTTATGCAACAACCTTTCCAAAAC |
| AV110_56-Fw | CAGGAGGAATATACCATGCTAAGTGAAAACCCAATTACAG |
| AV110_56-Rv | CACGACGCTGAGGTGTTATAGTGTTCCATATACTCGAGTT |
| AV110_60-Fw | CAGGAGGAATATACCATGACTAAAGAACAGCATG |
| AV110_60-Rv | CACGACGCTGAGGTGCTATTTTACTAAAGATTCAATATAC |
| AV110_61-Fw | CAGGAGGAATATACCATGATTAAATTAACTGTGGCTGT |
| AV110_61-Rv | CACGACGCTGAGGTGTCATTGGCATTTTTCCTCTTTT |
| AV110_63-Fw | CAGGAGGAATATACCATGAATATGCAATTGATTACTAACG |
| AV110_63-Rv | CACGACGCTGAGGTGTTACTTTAAGCATGTTTGAATAATC |
| AV110_72-Fw | CAGGAGGAATATACCATGCTTAAATTAGTATTCGCATG |
| AV110_72-Rv | CACGACGCTGAGGTGTCATTTATAAACGCTCTCAGTAATTTC |
| AV110_74-Fw | CAGGAGGAATATACCATGGAAATTGGAAAATCTTATATCA |
| AV110_74-Rv | CACGACGCTGAGGTGCTACTTTTTAAAAGTTTTTTGCA |
| AV110_92-Fw | CAGGAGGAATATACCATGAATCTTAAACAACTCCAG |
| AV110_92-Rv | CACGACGCTGAGGTGTCAATTATTCTCATCATCTAATTTC |
| AV110_93-Fw | CAGGAGGAATATACCATGAAAACTTTTAAAGAACGTTTAG |
| AV110_93-Rv | CACGACGCTGAGGTGTTAGTAAAGGTCCTCAGAGTAAAG |
| AV110_94-Fw | CAGGAGGAATATACCATGAATAAGCTGAGTATTATTAACG |
| AV110_94-Rv | CACGACGCTGAGGTGTCATTTAAACCATCCTTTAATACG |
| AV110_100-Fw | CAGGAGGAATATACCATGGTATCATGGATTATTGCATTGT |
| AV110_100-Rv | CACGACGCTGAGGTGTTACTCGCCTTTTAACACGTT |
| AV110_101-Fw | CAGGAGGAATATACCATGTCCAAATATTTAACTCGTAAAG |
| AV110_101-Rv | CACGACGCTGAGGTGTTAGTAACTCTGCAGAATGAATTTG |
| AV110_102-Fw | CAGGAGGAATATACCATGGCTAAGCTTTTTAAAGATGTTG |
| AV110_102-Rv | CACGACGCTGAGGTGTTATTCGGCTATATCAGAACG |
| AV110_108-Fw | CAGGAGGAATATACCATGATTATTGAAACGGCTAAAGAAAC |
| AV110_108-Rv | CACGACGCTGAGGTGTTATTTGGCCGCTTCCAC |

| | |
|---|---|
| AV110_109-Fw | CAGGAGGAATATACCATGAGAACATATAACGTGGACTTG |
| AV110_109-Rv | CACGACGCTGAGGTGTCACAAGTTACGTTTAAAAATTTC |
| AV110_114-Fw | CAGGAGGAATATACCATGGCTAATAAATTCCGCGTT |
| AV110_114-Rv | CACGACGCTGAGGTGTTATTGTTCCTTAAAGTGAGCTTTC |
| AV110_122-Fw | CAGGAGGAATATACCATGGTTTATGTATATGCAATAGTTT |
| AV110_122-Rv | CACGACGCTGAGGTGTCATCGTAAAGTCCCTGCA |
| AV110_135-Fw | CAGGAGGAATATACCATGGCACACTTTAACGAATG |
| AV110_135-Rv | CACGACGCTGAGGTGTTAATAACCTCGGTCTTGACG |
| AV110_140-Fw | CAGGAGGAATATACCATGATTTATTATATGCACAAAAATC |
| AV110_140-Rv | CACGACGCTGAGGTGTTAACCTCGATTCATAAATGC |
| AV110_143-Fw | CAGGAGGAATATACCATGAAAAAATTAATCGCTTTAGCA |
| AV110_143-Rv | CACGACGCTGAGGTGTTAATTAAATTTGTTTAAACGTTCA |
| AV110_168-Fw | CAGGAGGAATATACCATGACGAAGACTGATTATGAAATCC |
| AV110_168-Rv | CACGACGCTGAGGTGTCATATGTATTCTCTTCGAATGAA |
| AV110_178-Fw | CAGGAGGAATATACCATGTCTACTAAAATCAAAAACGTAG |
| AV110_178-RV | CACGACGCTGAGGTGTCATTTTACCGCCTTAACAATTTTC |
| AV110_180-Fw | CAGGAGGAATATACCATGAATCCTGAATCTATGTTATCGC |
| AV110_180-Rv | CACGACGCTGAGGTGTTAACCGAACTTTTTAGCGCC |
| AV110_181-Fw | CAGGAGGAATATACCATGAATATCAAACGAATGCTTTTT |
| AV110_181-Rv | CACGACGCTGAGGTGTCATTTTGTGTTCCACACCCCA |
| AV110_182-Fw | CAGGAGGAATATACCATGGGTAAAACATATCGTCGTAAAG |
| AV110_182-Rv | CACGACGCTGAGGTGTCAACTGTAACGATAACATTCG |
| AV110_184-Fw | CAGGAGGAATATACCATGACAATTGAAGATAAAGAAATTAAG |
| AV110_184-Rv | CACGACGCTGAGGTGTCATGGCTTAATTTCCTTCGC |
| AV110_188-Fw | CAGGAGGAATATACCATGTATAAATTTCGTAAAGGTTTAGC |
| AV110_188-Rv | CACGACGCTGAGGTGTTAAGCCTTTTTATCAAGAACAG |
| AV110_199-Fw | CAGGAGGAATATACCATGACACGTTCACATTACATTGA |
| AV110_199-Rv | CACGACGCTGAGGTGTTATAACACCTCAATGGCATTTC |
| AV110_201-Fw | CAGGAGGAATATACCATGAACGCTAAAGATATTTTCAACC |
| AV110_201-Rv | CACGACGCTGAGGTGTTATGCGTGAACCGTCTTTAAAC |
| AV110_202-Fw | CAGGAGGAATATACCATGCTCCAGTACGACAAG |
| AV110_202-Rv | CACGACGCTGAGGTGCTAAAAGAAACTTTTTTCGGTATTT |
| AV110_204-Fw | CAGGAGGAATATACCATGTTCAACGTTCAAATCAAAAAAG |
| AV110_204-Rv | CACGACGCTGAGGTGTTATTTCAGTAAAGTTACTTCAGAG |
| AV110_208-Fw | CAGGAGGAATATACCATGAAAACTGTAATTGAAACAACC |
| AV110_208-Rv | CACGACGCTGAGGTGTTATTTCAAAGTTGGGATAATAGAA |
| AV110_214-Fw | CAGGAGGAATATACCATGTCACGTAAAGAAAAAATTTCTAAG |
| AV110_214-Rv | CACGACGCTGAGGTGTCACATAATCTCGTTAACATACATT |
| AV110_224-Fw | CAGGAGGAATATACCATGACTATCAATTCAGACGTTT |
| AV110_224-Rv | CACGACGCTGAGGTGCTAGATTTTAATCACTGCTTTAGA |
| AV110_229-Fw | CAGGAGGAATATACCATGATGACCACTATTGAAGT |
| AV110_229-Rv | CACGACGCTGAGGTGTCAATTAGAAAATAAATTTATCCAAG |
| AV110_230-Fw | CAGGAGGAATATACCATGAATGCTTTTACACACCATTCTG |
| AV110_230-Rv | CACGACGCTGAGGTGTCATTGCATTCCAATCACAA |
| AV110_250-Fw | CAGGAGGAATATACCATGCCTATTAATTGGGTAGGATATTG |
| AV110_250-Rv | CACGACGCTGAGGTGTTAAGAGTTGTGGCTTTTGCATC |

| | |
|---|---|
| AV110_251-Fw | CAGGAGGAATATACCATGCGAAACCAAAATAATGATATTG |
| AV110_251-Rv | CACGACGCTGAGGTGTTACTTGGCATGTATAGGGTTGG |
| AV110_253-Fw | CAGGAGGAATATACCATGGCTAAAGTTAATATTGATGT |
| AV110_253-Rv | CACGACGCTGAGGTGTTATTTTTTAATCAAATTGACATAAAAC |
| AV110_255-Fw | CAGGAGGAATATACCATGAGTCTTTACACTGAACTG |
| AV110_255-Rv | CACGACGCTGAGGTGTCATTTATTTTTAATTTTGCAAATT |
| AV110_262-Fw | CAGGAGGAATATACCATGATTCTTTATGCTAAAGTAAAAT |
| AV110_262-Rv | CACGACGCTGAGGTGTCACCGTACCATATTAATGTAATT |
| AV110_263-Fw | CAGGAGGAATATACCATGAAATATCACGTATATGCGGAT |
| AV110_263-Rv | CACGACGCTGAGGTGTCATGAATGTTCCACCATTTCA |
| AV110_264-Fw | CAGGAGGAATATACCATGAAACGCGCTGAACTG |
| AV110_264-Rv | CACGACGCTGAGGTGTTACTCACCTTTTGCAATTTTATCC |
| Gp94-Halo_Rv | TTTAAACCATCCTTTAATACGTTGCC |
| Halotag_Rv | CACGACGCTGAGGTGTCAGCCGGAAATCTCGAGC |
| Halotag-gp94_Fw | AAAGGATGGTTTAAAGGAGGAGGTGGAAGTCC |
| pBTB3-lin_Fw | CCTCAGACTCCAGCGTAAC |
| pBTB3-lin_Rv | GGTATATCTCCTTCTTCGTTGACGAATTCTCTAGCC |
| pBTB3_Fw | ATTAGCGGATCCTACCTGAC |
| pBTB3_Rv | TATGTGGTCGAAGGCTGG |
| pBTB3-MreB_Fw | AGAAGGAGATATACCATGTTGAAAAAATTTCGTGGCATG |
| pBTB3-MreB_Rv | CGCTGGAGTCTGAGGTTACTCTTCGCTGAACAGGTC |
| pBTB3-MreC_Fw | AGAAGGAGATATACCATGAAGCCAATTTTTAGCCG |
| pBTB3-MreC_Rv | CGCTGGAGTCTGAGGACGATAGCTCGCCACTATTG |
| pBTB3-MreD_Fw | AGAAGGAGATATACCGTGGCGAGCTATCGTAG |
| pBTB3-MreD_Rv | CGCTGGAGTCTGAGGTTATTGCACTGCAAACTGCTG |
| pUT18-lin_Fw | CCGAGCTCGAATTCAGCCG |
| pUT18-lin_Rv | CTCTAGAGTCGACCTGCAGG |
| pUT18-insert_Fw | TTTATGCTTCCGGCTCGTATG |
| pUT18-insert_Rv | CTATGCGGCATCAGAGCA |
| pUT18_Gp94_Fw | AGGTCGACTCTAGAGATGAATAAGCTGAGTATTATTAACGAACTTCGTAAATG |
| pUT18_Gp94_Rv | TGAATTCGAGCTCGGTTTAAACCATCCTTTAATACGTTGCCAAATAGATTTTTGT |
| pUT18C_Gp94_Fw | AGGTCGACTCTAGAGATGAATAAGCTGAGTATTATTAACG |
| pUT18C_Gp94_Rv | TGAATTCGAGCTCGGTTTAAACCATCCTTTAATACG |
| pTK25-lin_Fw | CCTAAGTAAGTAAGAATTCACTGGC |
| pTK25-lin_Rv | CTCTAGAGTCGACCCTGCAG |
| pTK25-insert_Fw | TTATGCTTCCGGCTCGTATG |
| pTK25-insert_Rv | CGGCATCAGAGCAGATTGTA |
| pTK25_MreB_Fw | GGGTCGACTCTAGAGATGTTGAAAAAATTTCGTGGCATGTTTTCC |
| pTK25_MreB_Rv | TCTTACTTACTTAGGCTCTTCGCTGAACAGGTCGCC |
| pTK25_MreC-Fw | GGGTCGACTCTAGAGATGAAGCCAATTTTTAGCCGTG |
| pTK25_MreC-Rv | TCTTACTTACTTAGGTTGCCCTCCCGGCG |
| pTK25_MreD _Fw | GGGTCGACTCTAGAGGTGGCGAGCTATCGTAGCCA |
| pTK25_MreD _Rv | TCTTACTTACTTAGGTTGCACTGCAAACTGCTGACG |
| pTK25_FtsZ _Fw | GGGTCGACTCTAGAGATGTTTGAACCAATGGAACTTACCAATGAC |
| pTK25_FtsZ _Rv | TCTTACTTACTTAGGATCAGCTTGCTTACGCAGGAAT |
| pNTK25-lin_Fw | AGCTCGAATTCAATGACCATG |

| | |
|---|---|
| pNTK25-lin_Rv | CTCTAGAGTCGACCTGCAGG |
| pNTK25-insert_Fw | TTATGCTTCCGGCTCGTATG |
| pNTK25-insert_Rv | CGGCATCAGAGCAGATTGTA |
| pNTK25_MreB_Fw | AGGTCGACTCTAGAGATGTTGAAAAAATTTCGTGGCATGTTTTCC |
| pNTK25_MreB_Rv | CATTGAATTCGAGCTCTCTTCGCTGAACAGGTCGCC |
| pNTK25_MreC-Fw | AGGTCGACTCTAGAGATGAAGCCAATTTTTAGCCGTG |
| pNTK25_MreC-Rv | CATTGAATTCGAGCTTTGCCCTCCCGGCG |
| pNTK25_MreD _Fw | AGGTCGACTCTAGAGGTGGCGAGCTATCGTAGCCA |
| pNTK25_MreD _Rv | CATTGAATTCGAGCTTTGCACTGCAAACTGCTGACG |
| pNTK25_FtsZ _Fw | AGGTCGACTCTAGAGATGTTTGAACCAATGGAACTTACCAATGAC |
| pNTK25_FtsZ _Rv | CATTGAATTCGAGCTATCAGCTTGCTTACGCAGGAATG |

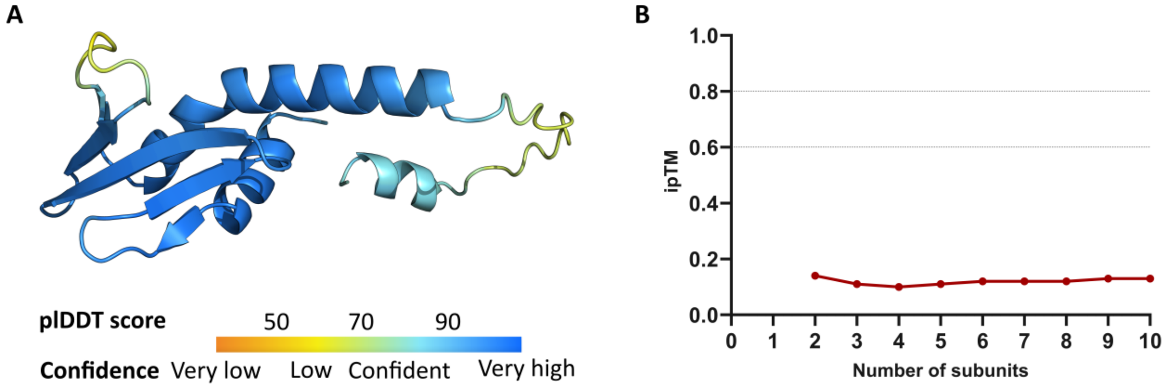


**Figure S1: Predicted protein structure and multimerization of Gp94, related to** Error! Reference source not found.**.** A) The protein structure of Gp94 was predicted with AlphaFold 3 and colored according to the pIDDT confidence score of each amino acid. B) Gp94 was predicted to function as a monomer based on the ipTM score of the AlphaFold 3 predictions of the different multimers.

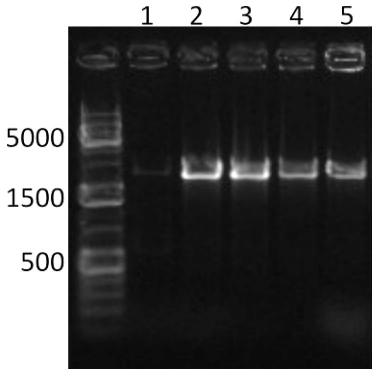


**Figure S2: Colonies obtain resistance against Gp94 due to an insertion in the plasmid, related to** Error! Reference source not found.**.** Colony PCR amplifying the promoter and insert region of the plasmid using five colonies that became resistant against Gp94 shows an insertion in this region for all tested colonies. The expected amplicon size of pPS26::*gp94* is 632 bp.

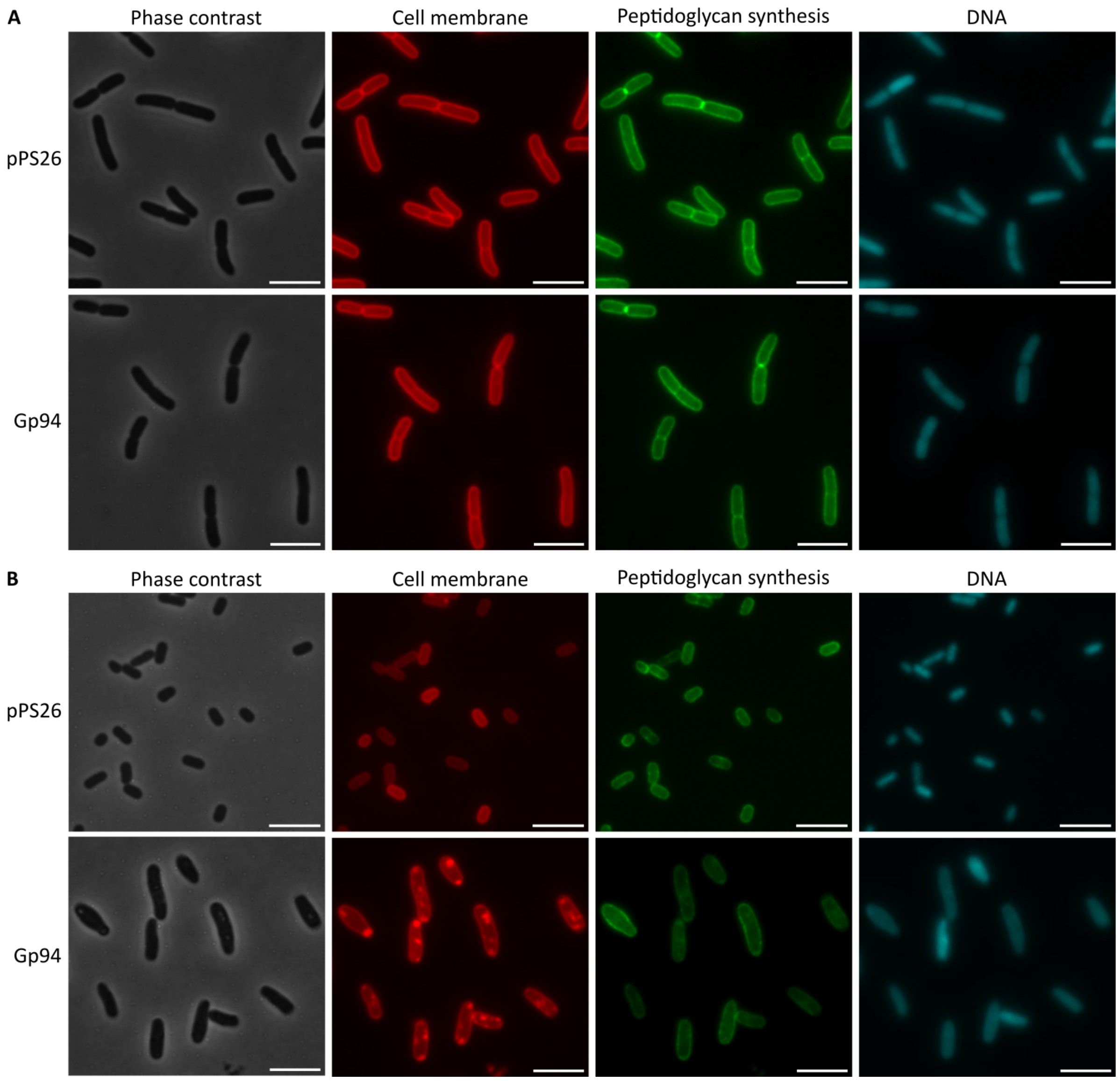


**Figure S3: Phase contrast and fluorescence microscopy after 1 and 4 h expression of Gp94, related to** Error! Reference source not found.**.** ESBL098 transformed with the empty vector or pPS26::*gp94* after A) 1 h and B) 4 h of induction with staining of the cell membrane, peptidoglycan synthesis, and the DNA. Scale bar = 5 µm.

**Video S1**: Time-lapse microscopy of Gp94 expression in ESBL098, related to **Error! Reference source not found.**. The growth of ESBL098 pPS26::*gp94* on an LA pad supplemented with rhamnose was followed for 25 h. Scale bar = 10 µm.

**Video S2**: Time-lapse microscopy of Gp94 expression in ESBL098 with live-dead staining, related to **Error! Reference source not found.**. The growth of ESBL098 pPS26::*gp94* on an LA pad supplemented with rhamnose and propidium iodide was followed for 27 h. Scale bar = 10 µm.
